# A Deep Learning Approach for Rapid Mutational Screening in Melanoma

**DOI:** 10.1101/610311

**Authors:** Randie H. Kim, Sofia Nomikou, Nicolas Coudray, George Jour, Zarmeena Dawood, Runyu Hong, Eduardo Esteva, Theodore Sakellaropoulos, Douglas Donnelly, Una Moran, Aristides Hatzimemos, Jeffrey S. Weber, Narges Razavian, Ioannis Aifantis, David Fenyo, Matija Snuderl, Richard Shapiro, Russell S. Berman, Iman Osman, Aristotelis Tsirigos

**Author notes:** These authors contributed equally to this work (Randie H. Kim, Sofia Nomikou). Corresponding Authors: Aristotelis Tsirigos, PhD, Associate Professor of Pathology, Director, Applied Bioinformatics Laboratories, NYU Grossman School of Medicine, 227 East 30^th^ Street, New York, New York, Iman Osman, MD, Professor of Dermatology, Associate Dean for Translational Research Support, Director, Interdisciplinary Melanoma Cooperative Group, NYU Grossman School of Medicine, 522 First Avenue.

## Abstract

Image-based analysis as a rapid method for mutation detection can be advantageous in research or clinical settings when tumor tissue is limited or unavailable for direct testing. Here, we applied a deep convolutional neural network (CNN) to whole slide images of melanomas from 256 patients and developed a fully automated model that first selects for tumor-rich areas (Area Under the Curve AUC=0.96) then predicts for the presence of mutated *BRAF* in our test set (AUC=0.72) Model performance was cross-validated on melanoma images from The Cancer Genome Atlas (AUC=0.75). We confirm that the mutated *BRAF* genotype is linked to phenotypic alterations at the level of the nucleus through saliency mapping and pathomics analysis, which reveal that cells with mutated *BRAF* exhibit larger and rounder nuclei. Not only do these findings provide additional insights on how *BRAF* mutations affects tumor structural characteristics, deep learning-based analysis of histopathology images have the potential to be integrated into higher order models for understanding tumor biology, developing biomarkers, and predicting clinical outcomes.

## Introduction

Mutations in the *BRAF* oncogene are found in 50-60% of all melanomas^1^. With the development of targeted therapies^2, 3^, determining the mutational status of *BRAF* has become an integral component for the management of Stage III/IV melanomas. Current methods for mutation detection include DNA molecular assays^4^ and rapid screening tests, such as immunohistochemistry, real-time polymerase chain reaction (PCR) and automated platforms^5, 6, 7^, all of which require tumor tissue for analysis. Recently, image-based analysis has been investigated as an alternative method for mutation prediction, which can be particularly useful in settings when tumor is either not available or inadequate for direct testing. While many of these studies involve the use of radiomics^8^, image-based analysis has expanded to histopathology with the advent of digitized whole slide images (WSI).

The field of pathomics attempts to extract and quantitate features from high-resolution digitized WSI on a large scale for the purposes of integrating with molecular signatures, developing biomarkers, and predicting clinical or treatment outcomes^9^. These tasks include quantifying the number of objects, detecting object boundaries, classifying groups of objects, and labeling that allow for characterization of tissue not typically possible by traditional microscopic evaluation^10^. With the amount of data that can be potentially generated with pathomics, machine learning algorithms are uniquely positioned to link image features to a greater framework of understanding tumor biology^11, 12^.

Relatedly, deep convolutional neural networks (CNN) have been shown to predict for the presence of actionable genetic mutations, such as *EGFR, ER,* and *BRAF* in a number of solid tumors using histopathological images^13, 14, 15, 16^, demonstrating that genotypic-phenotypic changes can be detected in tumor cells and/or the tumor microenvironment. In response to limitations that deep learning algorithms represent a “black box”, additional studies have attempted to correlate learned histopathologic features with specific phenotypes^17^. Furthermore, better understanding of how various training parameters and modes of learning can influence model performance is required before broader applications to clinical practice.

In this study, we utilize two distinct and complementary methods of analyzing whole slide images for the prediction of mutated *BRAF* in melanomas resected from patients prospectively enrolled in a single-institution, IRB-approved clinicopathological biorepository. First, we apply deep learning techniques to histopathology images of FFPE primary melanomas in order to develop a model from tissue specimens that are more representative of what might be seen in routine clinical practice. Through saliency mapping, we determine that cell nuclei are a key feature in what our network learns for mutation prediction. Finally, we confirm that the mutated *BRAF* genotype is associated with detectable and quantifiable nuclear differences using pathomics analysis, thus providing a genotype-phenotype link in melanoma tumor cells. We present our deep learning models for predicting *BRAF* mutations in melanoma to demonstrate the feasibility and explainability of rapid image-based mutational screening that can be used in research or clinical-based settings in which limited tumor tissue is available for direct testing.

## Results

### Dataset characteristics

#### NYU cohort

Formalin-fixed paraffin embedded (FFPE) hematoxylin and eosin (H&E)-stained slides of 293 primary melanomas from 256 unique patients were included in this study. 103 melanomas harbored mutated *BRAF* and 190 melanomas were wild-type *BRAF.* All slides were digitized at 20x magnification and reviewed for quality control. Images that were blurry, faded, or contained no tumor were excluded. Additionally, only the slide with the greatest tumor content was used to build the classifier in order to reduce bias, leading to a final data cohort of 256 H&E slides. Slides were divided into training (n=184), validation (n=36), and independent testing cohorts (n=36) without overlap between patient subsets. Within each cohort, *BRAF*-mutant and *BRAF-* wild type (*BRAF-WT*) melanomas were represented. V600E comprised 70% of the *BRAF* mutations.

#### The Cancer Genome Atlas (TCGA) cohort

An image dataset of 68 digitized FFPE H&E-stained slides of primary melanomas^18^ were retrieved from TCGA database^19^ and used as a second independent cohort. Clinical information was not available for all slides. Because TCGA primary melanoma specimens are enriched for thicker tumors (median=2.7mm; mean=4.9mm^18^), we selected 28 specimens with Breslow depth similar to our cohort as a second independent cohort to maintain uniformity of Breslow depth in our analysis.

### Automated selection of primary melanomas on whole slide histopathology images

Our computational workflow is shown in **Figure 1A** and is the same across all our classifiers (see Methods). Because skin excisions often contain heterogeneous tissue, our first task was to automate the identification of melanoma on whole slide images. Tumor-rich areas were manually annotated “in” the regions of interest (ROI) by a single dermatopathologist while normal skin, associated appendages, connective and subcutaneous tissue, necrosis, hemorrhage, and aggregates of dense inflammation were “out” of the ROI. For this task, we chose the Inception v3 architecture, which has been previously shown to accurately distinguish between tumor and non-tumor areas on H&E slides^13^. Learning curves are presented in **Figure 1B left and middle**. Model performance achieved a per patient AUC=0.96 [95% CI: 0.90-0.99] and a per tile AUC=0.92 [95% CI: 0.918-0.921] (**Figure 1B right**). H&E-stained non-annotated whole slides of *BRAF*-mutant and *BRAF-WT* melanomas along with their corresponding network-generated probability heat maps and pathologist-annotated tumor masks are presented in **Figure 1C.** Notably, there is excellent concordance between the pathologist and the network. Training performed on images at 10x and 5x magnification resulted in similar network performances (**Supplemental Figure 1** and **Supplemental Table 1**). The networks generated by this analysis are hereafter referred to as “TumorNet” along with the corresponding magnification.

**Figure 1.**
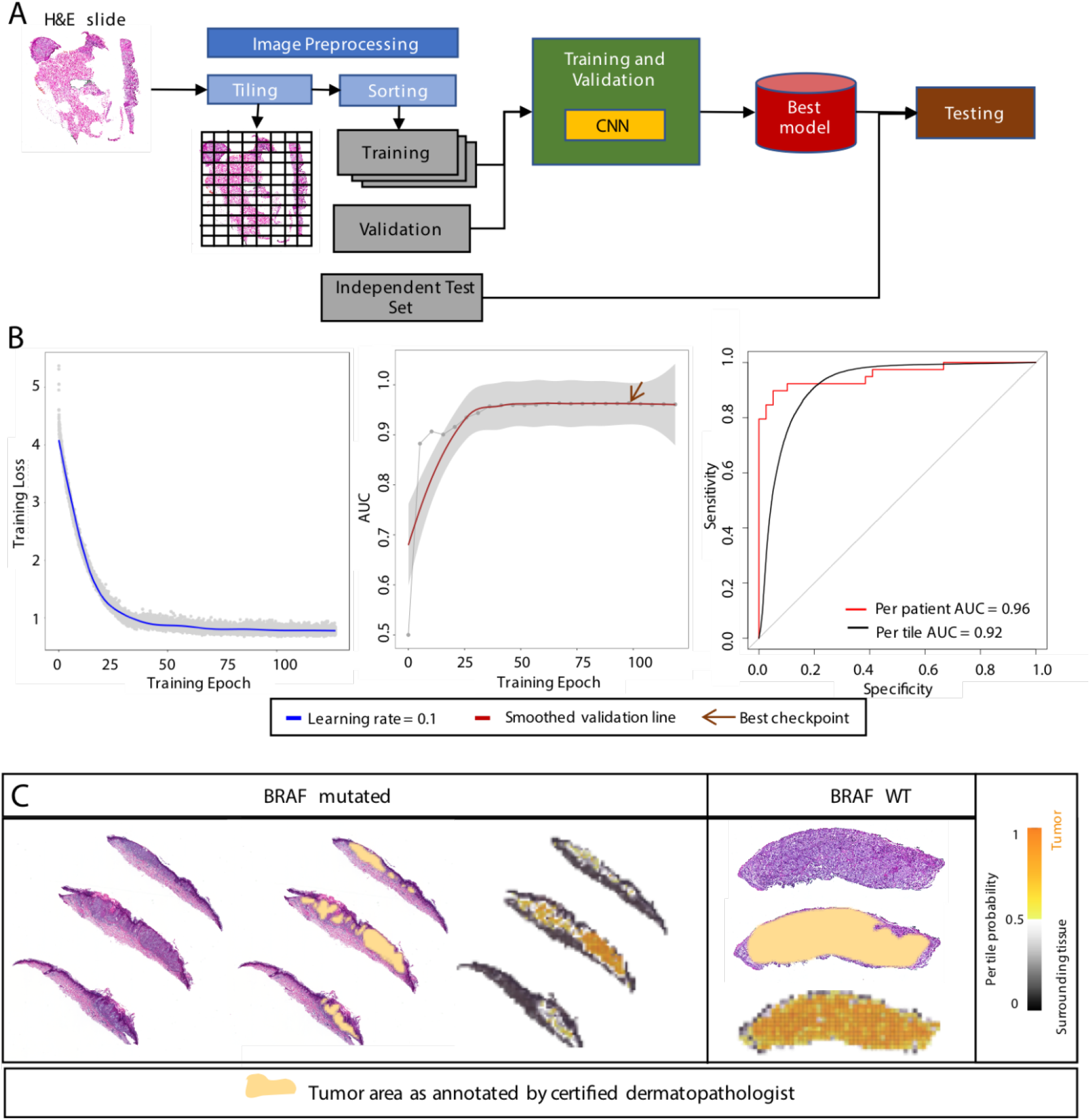
Automated tumor annotation. **A**. Computational workflow for all our classifiers. To train the CNN architectures, slides are tiled to non-overlapping tiles and assigned to training, validation and independent sets comprising of 70%, 15% and 15% of the total number of tiles, respectively. After conversion to TF Record format, training is performed. The best performing model on the validation data is evaluated on the independent set. **B.** Training loss (left) of Inception v3 for tumor annotation. Validation AUC (middle) across training with best model chosen at 98 training epochs. ROC curves on test set per tile and per patient (right). **C.** Examples of a *BRAF* mutated and a *BRAF* WT slide for the tumor annotation classifier with corresponding tumor areas as annotated by certified dermatopathologist.

### BRAF mutation prediction from melanoma whole-slide images by different CNN architectures

We first decided to explore the performance of three state-of-the-art CNN architectures in *BRAF* mutation prediction; Inception v3, VGG16^20^ and ResNet18^21^. All three architectures were successfully trained from scratch on the same dataset split into training, validation and test sets (Panels a and b of **Figures 2A-C**). Performance on the independent test set was varied, with Inception v3 achieving an AUC=0.69 [95% CI:0.50-0.86] (**Figure 2A** panel c); VGG16 achieving AUC=0.74 [95% CI:0.58-0.90] (**Figure 2B** panel c); and ResNet18 achieving AUC=0.86 [95% CI:0.74,0.99] (**Figure 2C** panel c). When applied to the TCGA dataset, Inception v3 generalized better (AUC=0.73 [95% CI:0.53-0.94] compared to AUC=0.59 [95% CI:0.37-0.82] and AUC=0.59 [95% CI:0.36-0.81] of VGG16 and ResNet18 respectively (Panel d of **Figures 2A-C** and **Supplemental Table 2**) (See Methods for details). Consequently, we chose Inception v3 as the most suitable architecture for our subsequent analyses.

**Figure 2.**
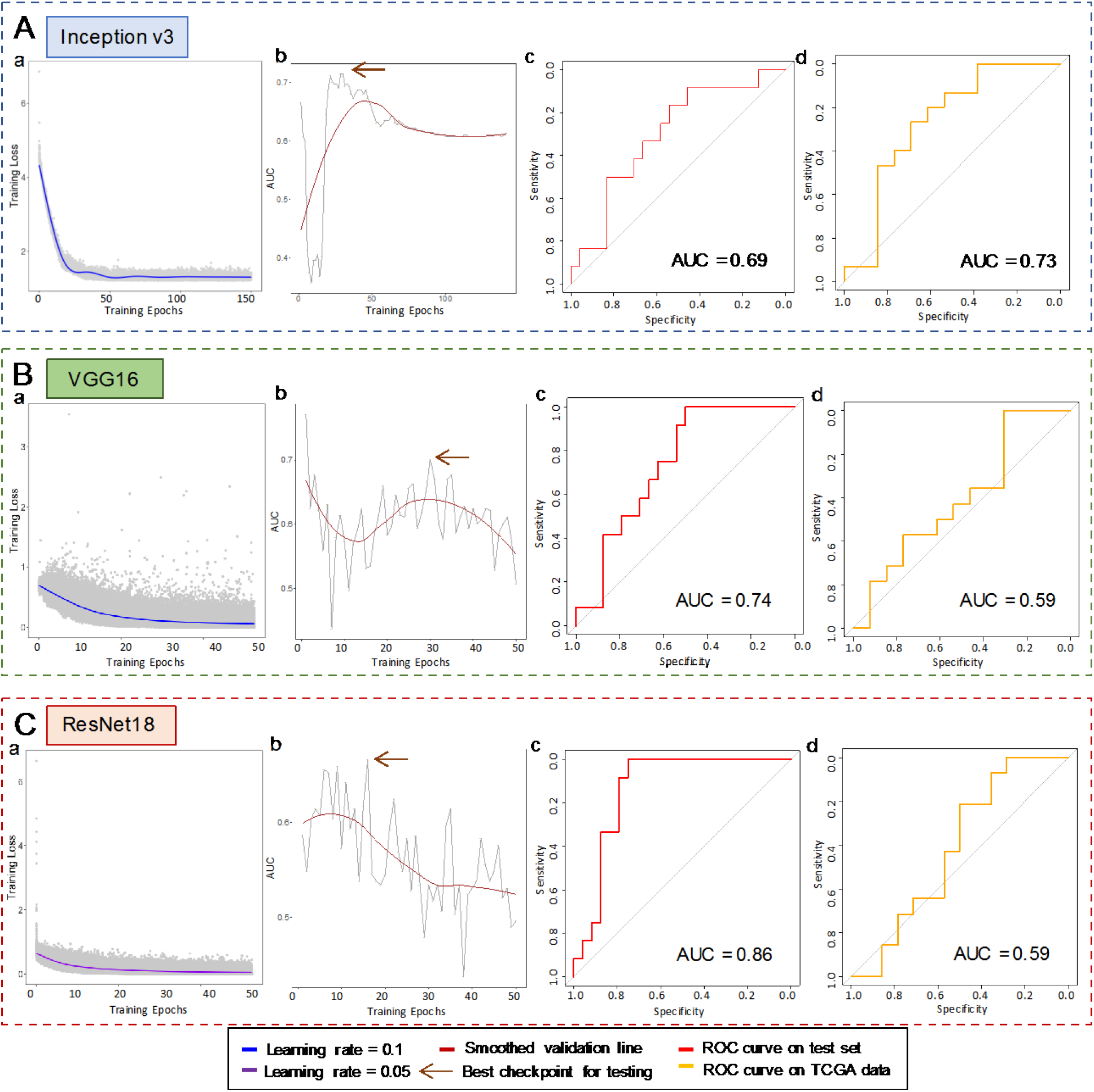
*BRAF* mutation prediction is feasible across multiple CNN architectures. **A)** Inception v3 a) Training Loss of Inception v3 for *BRAF* mutation prediction. b) Validation AUC across training. Best checkpoint is chosen at 30 training epochs. c) ROC curve for independent test set on best checkpoint. AUC is 0.69. d) ROC curve for external TCGA cohort on best checkpoint. AUC is 0.73. **B)** VGG16 a) Training Loss of VGG16 for *BRAF* mutation prediction. b) Validation AUC across training. Best checkpoint is chosen at 30 training epochs. c) ROC curve for independent test set on best checkpoint. AUC is 0.74. d) ROC curve for external TCGA cohort on best checkpoint. AUC is 0.59. **C)** ResNet18 a) Training Loss of ResNet18 for *BRAF* mutation prediction. b) Validation AUC across training. Best checkpoint is chosen at 16 training epochs. c) ROC curve for independent test set on best checkpoint. AUC is 0.86. d) ROC curve for external TCGA cohort on best checkpoint. AUC is 0.59.

### Effect of training parameters on BRAF mutation prediction using Inception v3

We next sought to elucidate the effect of tile size and training mode of Inception v3 on BRAF mutation prediction. Because Inception v3 only accepts tile sizes of 299×299 pixels, we used different magnifications as a proxy and retrained the architecture at 5x and 10x magnifications using the same data set split to training, validation and independent test sets. Additionally, we explored whether utilization of transfer learning to fine tune the last layer of the network influenced architecture performance compared to training all layers from scratch. For transfer training, we retrained the architecture using the weights of the ImageNet challenge^22^ as well as the weights of the best checkpoints from our own melanoma annotation classifiers for each magnification (see Methods for details). The networks’ performance on the independent test set and the TCGA cohort are shown in **Figure 3A** with additional details provided in **Supplemental Figures 2,3,4** and **Supplemental Table 3**. Training at 5x magnification yielded inconsistent results, with large variations in the AUC values. While training at 10x magnification performed more consistently across different training modes, training at 20x magnification demonstrated the least amount of variation, with the model trained with transfer training based on the weights from our TumorNet network achieving the best AUCs for the independent NYU test set (AUC = 0.72 [95% CI:0.53-0.87]) and the TCGA cohort (AUC = 0.75 [95% CI:0.57-0.94]) (**Figure 3B**). Examples of *BRAF*-mutant and *BRAF-WT* H&E–stained slides from the independent test set and TCGA cohorts are shown in **Figure 3C** along with their probability heat maps.

**Figure 3.**
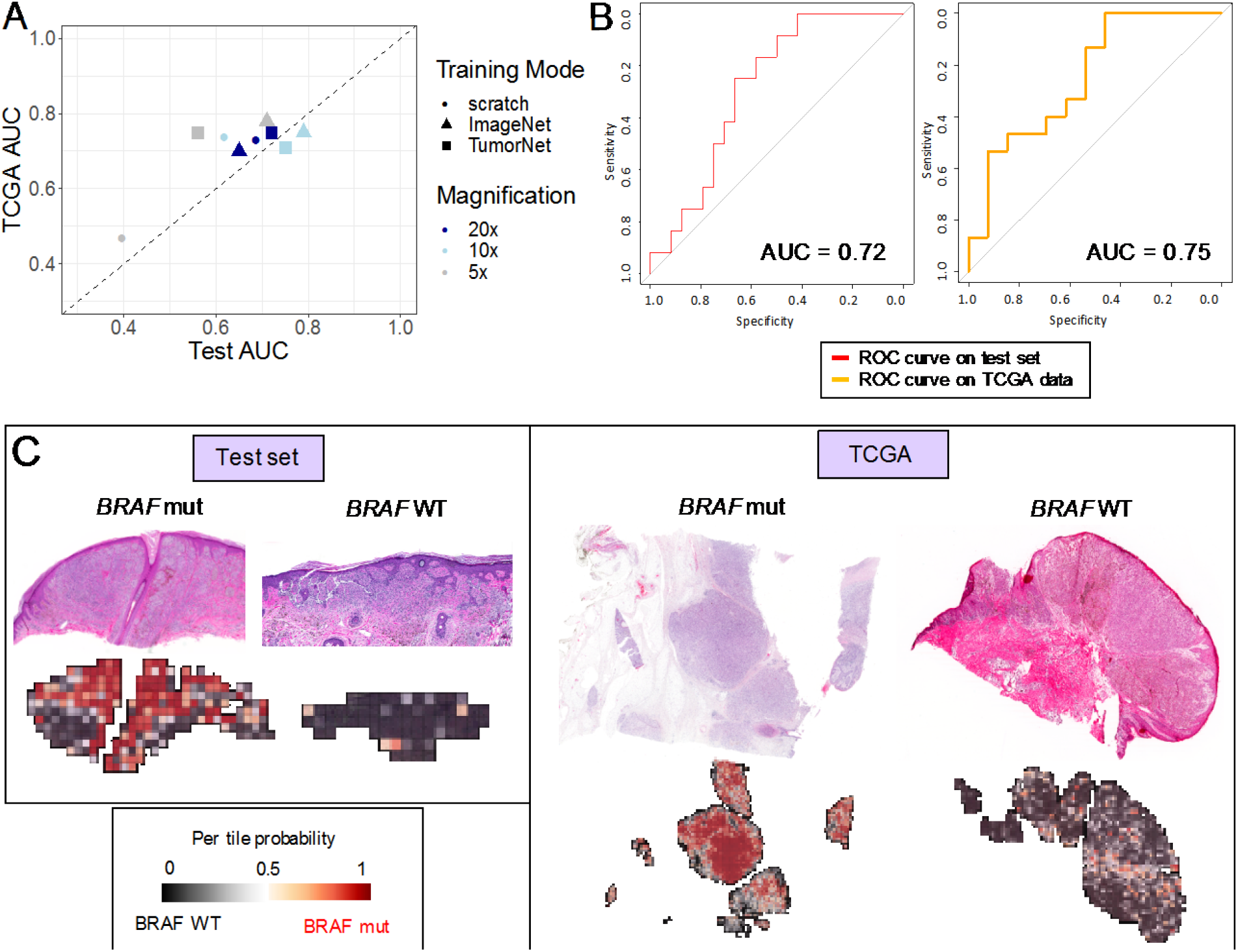
Exploration of the effect of magnification and learning mode on *BRAF* mutation prediction using Inception v3. **A.** Parameter exploration for magnification and training modes for Inception v3. The AUC on the independent test set and the external TCGA cohort are used as measures for prediction performance. Training at 5x seems unstable across different training modes (grey points). Training at 10x (light blue) and 20x (dark blue) yield more consistent results for different training approaches with 20x producing results with the smallest variation. Transfer training at 20x using the pre-trained tumor annotation network will be used onwards as our best classifier. **B.** ROC curve on independent test set for best performing checkpoint for classifier trained on tumor annotation network at 20x magnification. AUC is calculated at 0.72 CI[0.53-0.87] (left). ROC for external TCGA cohort with AUC at 0.75 CI[0.57-0.94]. **C.** Example mutation heat maps for *BRAF* mutated and *BRAF* WT slide from the test set (left) and the TCGA cohort (right). Tiles are colored based on their *BRAF* mutation probability values as predicted by the network.

Lastly, we investigated the effect of dataset size on prediction AUC. We down-sampled the dataset to 20,40,60 and 80% of initial data (**Supplemental Table 4**). Transfer training of Inception v3 using the weights of TumorNet20x was repeated for each down-sampled data set. Average AUC on the validation and test sets was reduced, as expected (**Supplemental Figure 5**). Fitting an inverse power law curve to the data demonstrated that in order for the classifier to achieve an AUC of 0.8, ~4.5x more data (i.e., at least 800 slides) would be needed. For the classifier to predict BRAF mutation with an AUC of 0.90, 10x more data (i.e. at least 1800 slides) would be needed.

### Automated sequential workflow for melanoma selection and mutation prediction

In order to improve utilization of our deep learning models, we developed a fully automated workflow by combining our tumor annotation and *BRAF* mutation prediction classifiers (**Figure 4**). For this task, we first verified that the BRAF mutation classifier trained on automatically annotated tumor areas performed similarly to the one trained on the manually annotated tumors. All 256 whole slide images (WSI) at 20x magnification were passed through the trained tumor annotation network (TumorNet). Tiles assigned with a probability of containing tumor higher than the threshold set were filtered and split into training, validation, and independent test sets. The Inception v3 architecture was re-trained on tiles selected by the automated network for mutation prediction. The network trained on tiles selected by TumorNet achieved similar performance to the one trained on the manually selected regions (**Supplemental Figure 6**), demonstrating a successful fully automated sequential model.

**Figure 4.**
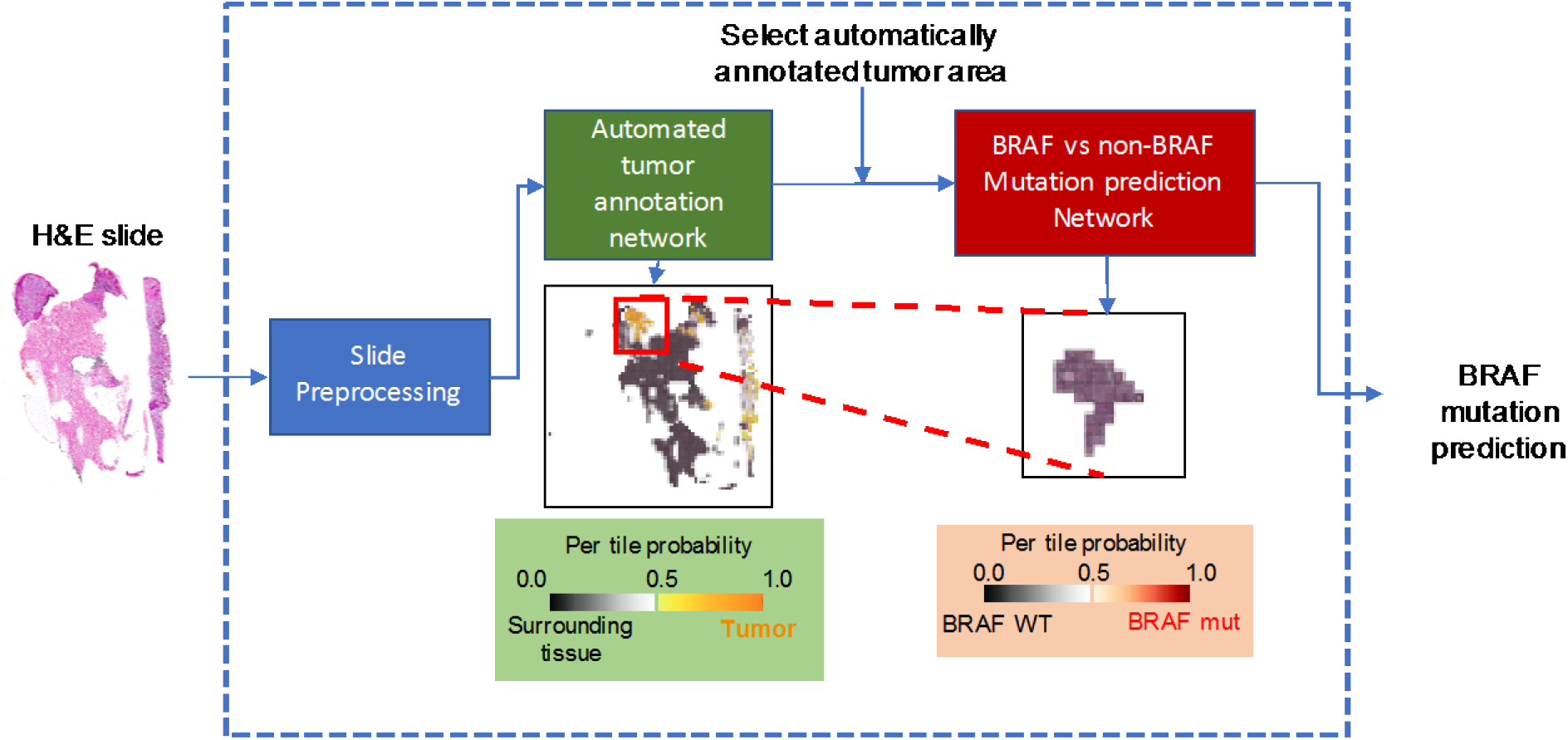
Sequential workflow for *BRAF* mutation prediction. Non-annotated whole slides are processed, tiled, and passed through the automated tumor annotation network which assigns a probability to each tile of belonging in the tumor. Tiles with high probability of containing tumor are subsequently passed through the mutation prediction network for determining the mutational status of the slide of interest.

### Association of network mutation localization with immunohistochemical analysis

To further corroborate network accuracy, we examined whether network-generated probability heat maps are true visual representations of mutation localization. An additional set of 17 *BRAF^V600E^* cases underwent automated algorithmic mutation prediction and immunohistochemical (IHC) analysis with the monoclonal VE1 antibody, a reliable screening tool for detecting the specific V600E mutation^23^. A single dermatopathologist blinded to mutational status manually annotated tumor ROI on H&E-stained slides as well as regions of positive staining on both the H&E-stained and IHC slides. (**Figure 5A**). The annotated mask of positive IHC staining and the mask for the annotated tumor area form the H&E slide were then overlaid on the network-generated probability heat map. The average probability of tiles falling inside vs. outside the selected antibody stained mask was calculated and is displayed in the form of box plots in **Figure 5B** for all 17 slides. From the 7 slides that were correctly predicted as BRAF mutant by the network, five of them show statistically significant higher BRAF probabilities for the tiles inside the annotated V600E antibody stained area compared to the remaining tumor tiles, indicating that the network indeed localizes mutated *BRAF.*

**Figure 5.**
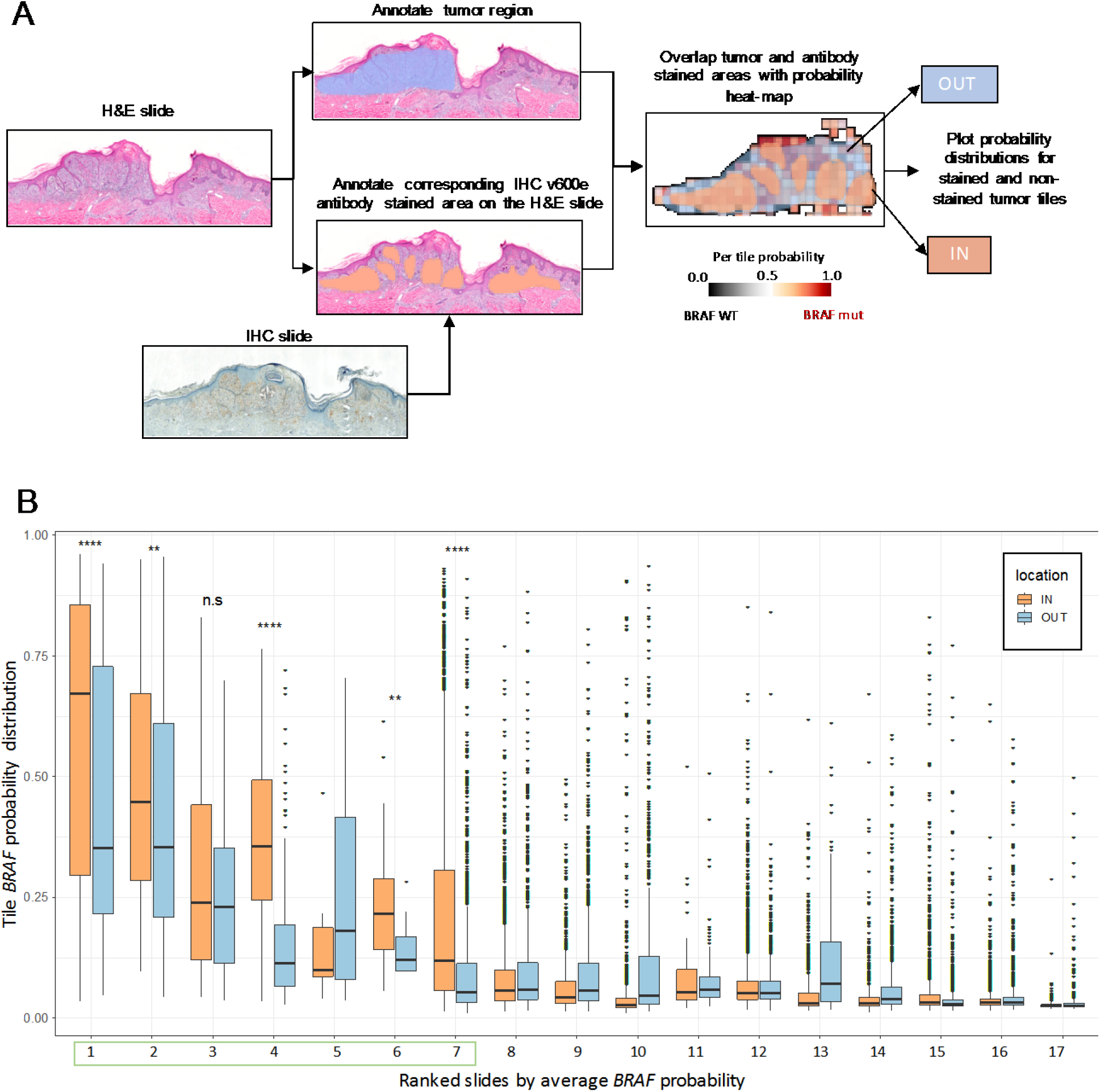
BRAF V600E-predicted tumor areas overlap with immunohistochemical V600E antibody staining for correctly predicted slides. **A)** Overlap strategy for IHC and H&E slides. Tumor annotation was performed on the H&E slides. Using the corresponding stained IHC slide for V600E, a single pathologist performed annotation of the respective area on the H&E slide to avoid potential inconsistencies due to the use of different slides to perform H&E and IHC if a different overlap approach was utilized. Then, the masks for the annotated areas are overlapped with the tile BRAF mutation probability heat-map to perform the overlap analysis. **B)** Probability distributions for 17 *BRAF* V600E slides for tiles inside and outside of the V600E stained areas. From the seven slides correctly predicted as *BRAF* V600E (green box), five of them show statistically significant higher *BRAF* probabilities for the tiles inside the annotated V600E antibody stained area compared to the remaining tumor tiles.

### Cell nuclei are informative areas for BRAF mutation prediction

We next attempted to delineate some of the learned image features that contribute to *BRAF* mutation prediction by the CNN. Tiles from the NYU independent test set were ranked by *BRAF* mutation probability. The top 100 and bottom 100 tiles were then used to create saliency maps using our best performing network (Inception v3 trained at 20x on the pre-trained tumor annotation weights). Saliency maps are generated using the weights of the last layer of the network before the fully connected layer. The map visualizes the importance of each image pixel for the prediction (see Methods for implementation details). **Figure 6A** demonstrates examples from high confidence and low confidence tiles from six different patients, in which the H&E tile containing tumor is shown on the left, the saliency map is shown in the middle, and the overlap of the two is shown on the right. In the saliency map, pixels assigned colors in the “warm” spectrum are considered important for mutation prediction while pixels assigned “cool” colors contribute less to the prediction. In both the high and low *BRAF*-mutant probability tiles, pixels with the highest contribution to the network performance are those corresponding to cell nuclei. The same analysis was repeated on tiles from the TCGA slides (**Figure 6B**) and again demonstrate that areas corresponding to cell nuclei seem to be the most important structures for the network’s prediction.

**Figure 6.**
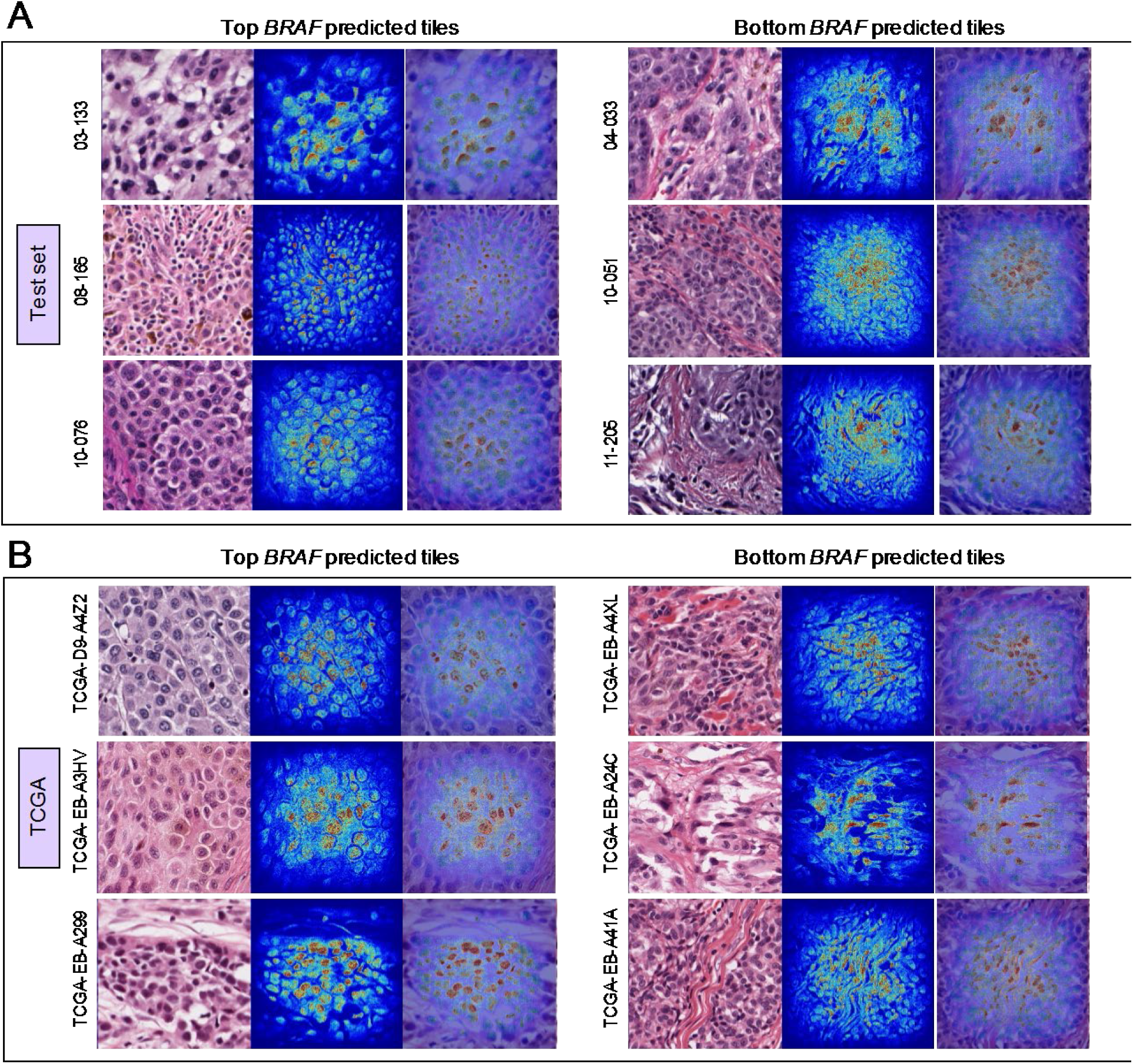
Saliency maps reveal cell nuclei as informative areas for *BRAF* mutation prediction. **A)** Saliency maps for three tiles predicted with highest *BRAF* probability (left) and three tiles predicted with the lowest *BRAF* probability (right) from six different patients in the independent NYU test set. **B)** Saliency maps for three tiles predicted with highest *BRAF* probability (left) and three tiles predicted with the lowest *BRAF* probability (right) from six different patients in the TCGA data set. It can be observed that for all tiles independently of the *BRAF* probability, the network considers cell nuclei to be the most informative structures for the prediction.

### Pathomics analysis reveals nuclear differences correlate to BRAF mutational status

To explore the feasibility of *BRAF* mutation prediction using traditional image analysis approaches we developed a Pathomics pipeline using CellProfiler, a publicly available software offering multiple functionalities for traditional image processing such as automated annotation of image structures^24^, to detect nuclei of tumor melanocytes (**Figure 7A**). Our pipeline focuses on annotating cells and nuclei from the H&E slide (see Methods for details). Our first task was to unmix colors that are present in H&E-stained slides, where hematoxylin stains nuclei blue-black and eosin stains proteins in the cytoplasm and connective tissue elements pink. Additionally, melanin pigment appears as brown granules. These color signals were de-convoluted to generate grayscale images that indicate the location of each stain with a white color. Because cells with high melanin content may represent melanophages rather than tumor melanocytes, the pigment channel was overlaid with the hematoxylin channel to identify highly pigmented cells. These cells were then removed from subsequent analysis (**Figure 7A**). Objects that passed criteria were measured and assessed for 18 features (**Supplemental Figure 7**, **8, 9 and 10**).

**Figure 7.**
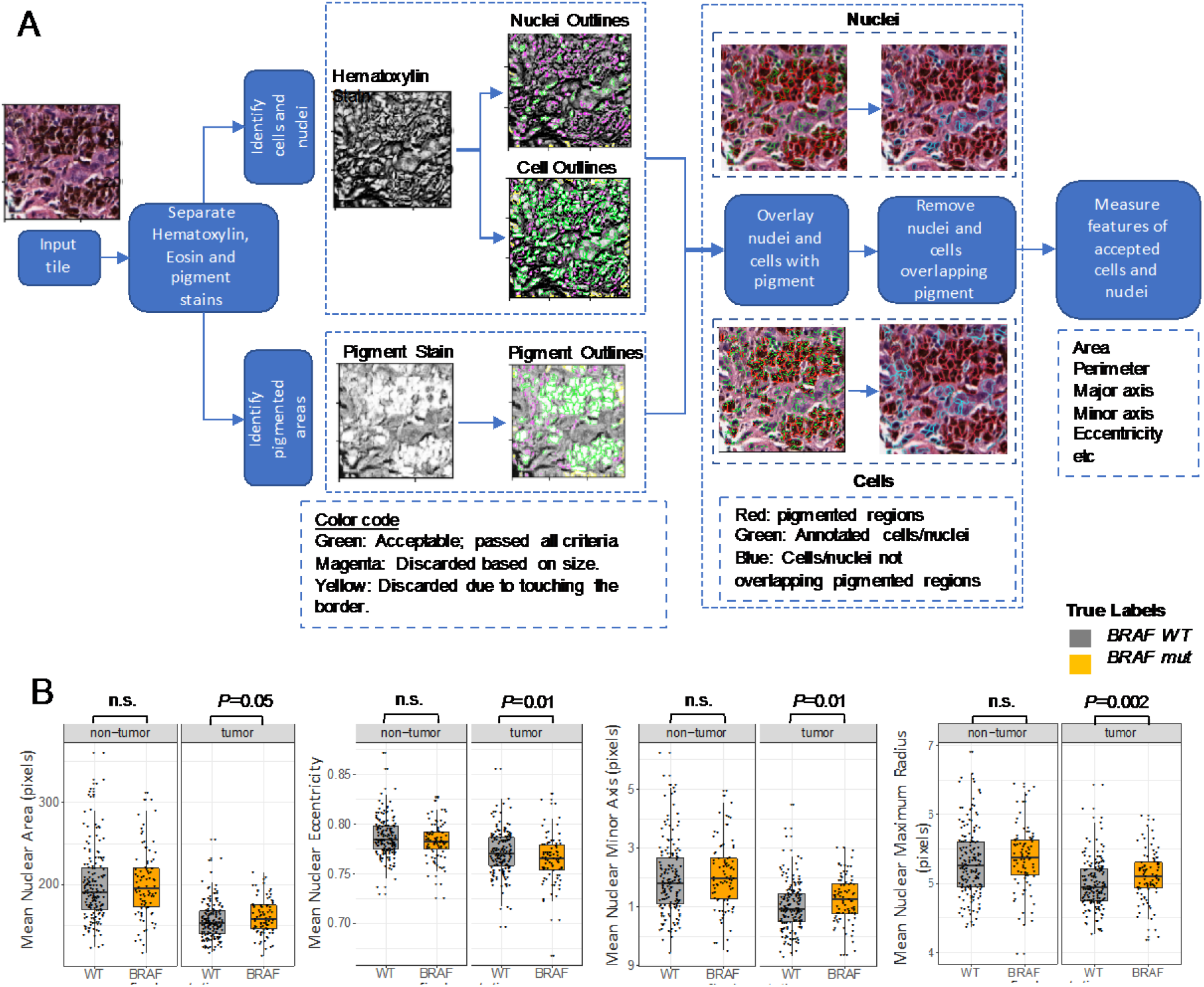
Pathomics analysis reveals that nuclear differences correlate to BRAF mutational status. **A)** Pathomics workflow with cellProfiler software. First, the hematoxylin, eosin and pigment stains are deconvolved. Hematoxylin is then used to annotate cells and cell nuclei. The pigment channel is used to annotate pigmented areas. Annotated nuclei and cells are overlapped with the pigmented regions and those overlapping the pigment are not considered for analysis. A variety of metrics for the size and shape of annotated nuclei are calculated and collected. **B)** Nuclear features for non-tumor and tumor nuclei aggregated per patient are plotted (for full list of features see Supplemental Figures 7,8,9 and 10). For non-tumor tiles, there are no differences between *BRAF* mutated and *BRAF* WT nuclei. For tumor tiles, *BRAF* mutated nuclei seem to have larger nuclear area, maximum radius and minor axis and lower eccentricity values. These results indicate bigger and rounder *BRAF* mutated nuclei compared to *BRAF* WT ones.

All 293 slides from our patient cohort were passed through the pipeline, which runs for each tile of a slide. The data were averaged across all identified nuclei and normalized by the total number of tiles by patient, when necessary. The analysis was performed on both tumor and non-tumor tiles of each slide. In non-tumor areas, there were no statistically significant differences in nuclear features across all melanomas (**Supplemental Figure 7**). In contrast, within tumor areas, differences in some nuclear features were detected between *BRAF*-mutated and *BRAF-WT* tumor tiles (**Figure 7B** and **Supplemental Figure 7**). These features included: (1) average nuclear area, (2) average nuclear eccentricity, (3) average minor axis, (4) average maximum, (5) median and (6) mean nuclear radius.

Compared to *BRAF-WT* nuclei, nuclei harboring mutated *BRAF* exhibited a larger average nuclear area with longer maximum, median, and mean nuclear radius, indicating that these nuclei are larger. Furthermore, *BRAF*-mutated nuclei demonstrated a longer minor axis and a smaller average nuclear eccentricity, indicating that the shape of the nucleus is rounder. The analysis was repeated on 64 available TCGA FFPE samples (**Supplemental Figure 8**) and demonstrated similar trends in nuclear features, although the differences did not reach statistical significance due to the small sample size. We also modified the pipeline to annotate and analyze cells instead of nuclei for both the NYU and the TCGA cohort (see Methods). No cellular features showed statistically significant differences across BRAF mutant and BRAF WT patients for the non-tumor tiles. In tumor tiles, the average cellular area, minor cell axis length and the maximum, median and mean cellular radii were larger in *BRAF*-mutant compared to *BRAF-WT* tumor nuclei for the NYU cohort. No significant differences were observed between *BRAF*-mutant and *BRAF-WT* tumor cells in TCGA data (**Supplemental Figures 9 and 10**).

Finally, we decided to explore if conventional pathomics image analysis can predict the BRAF mutation as well as our deep learning network. We trained a random forest model and a generalized linear model using 7-fold cross validation to mimic the number of ~37 slides in the independent test set that we have for our network. We used all our 293 slides and the 18 nuclear features provided by CellProfiler (**Supplemental Table 5** and Methods). The random forest model achieves an average AUC of 0.58 on the test set and 0.61 on the TCGA dataset. The generalized linear model yields an AUC of 0.56 on the test set and 0.58 on the TCGA data. Thus, deep learning was consequently better at predicting *BRAF* mutational status from H&E slides than conventional pathomics, an observation that has also been reported in radiomics studies^8^.

## Discussion

In the era of personalized medicine, molecular profiling can guide optimal cancer treatment, particularly if targeted therapies, such as *BRAF* inhibitors, are available. Predicting *BRAF* mutational status from image-based analysis is being investigated as an appealing method for rapid screening without the need for tumor tissue, and has been previously demonstrated in radiomics using ultrasound images for papillary thyroid cancer^25, 26^ and brain MRI images of metastatic melanomas^27^. More recently, deep CNN algorithms have been applied to histopathology images obtained from TCGA to predict for actionable mutations in lung adenocarcinoma^13^, papillary thyroid cancers^15^, and colorectal cancers^16^, indicating that genotypic alterations lead to phenotypic changes on the tumor cell level. In our study, we corroborate that *BRAF* mutations lead to specific morphologic changes, specifically larger and rounder nuclei, that can be predicted through deep learning and pathomics.

In melanoma, image-based analysis using deep learning has successfully been applied to classify pigmented lesions as benign vs. malignant using clinical^28^ or dermoscopic^29^ images with impressive accuracy. With respect to *BRAF* mutations, specific morphologic signatures associated with mutated *BRAF* in melanoma have been described independently with dermoscopy^30^, reflectance confocal microscopy^31^, and histology^32, 33^. These histologic features were determined by traditional microscopy and include greater pagetoid scatter, intraepidermal nesting, epidermal thickening, better circumscription, larger epithelioid and more pigmented melanocytes, and less solar elastosis. However, attempts to develop binary decision trees to predict for the *BRAF* mutation using histology alone achieved a predictive accuracy of only 60.3%^33^.

A pan-cancer deep learning image analysis by Kather et al.^16^ of FFPE H&E-stained slides of 14 different solid tumors and more than 5,000 patients from the TCGA database, successfully predicted for mutated *BRAF* in colorectal cancers. Interestingly, no significant mutations were able to be predicted from primary melanomas, and only *FBXW7* and *PIK3CA* from metastatic melanoma samples. One potential reason that mutation prediction was less successful in melanoma samples from TCGA data is the relatively small sample size. Here, we use a dataset of melanomas from over 250 patients to train our architecture, with our best model achieving an AUC=0.72. Importantly, we were able to cross-validate our model on images from TCGA [AUC=0.75]. We further substantiate the accuracy of our model by utilizing IHC analysis with the monoclonal VE1 antibody and assessing the overlay between positive IHC staining of BRAF^V600E^ on tissue sections and network-generated probability heat maps. Of the concordant cases between IHC and the network, 70% demonstrate significant overlap between the positive IHC staining and the heat map.

Despite the potential applications of unsupervised machine learning in pathology, a common concern is the “black box” issue in which learned features cannot be discovered from outputs. Relevant features can be inferred by examining high confidence image tiles for common morphological features. In the study by Kather et al.^16^, tiles of colorectal cancers ranked highly for mutated *BRAF* demonstrated areas of mucin as well as poorly differentiated tumor. A different computational approach by Fu et al.^17^ trained on 17,355 H&E-stained fresh-frozen tissue spanning 28 tumor types from TCGA to extract 1,536 image features and then used transfer learning to build prediction models for genotypes of interest. One high performing model was the association of mutated *BRAF* in papillary thyroid cancers, in which *BRAF* mutations are said to be found in 50%. The authors raise the question of whether the mutated *BRAF* genotype leads to the histological phenotype or whether *BRAF* mutations preferentially occur in certain cell types. We would argue the case for the latter, as *BRAF* mutations are also found in up to 50% of melanomas, but cannot be reliably predicted based on World Health Organization (WHO) histologic subtypes: superficial spreading melanoma, nodular melanoma, lentigo maligna melanoma, and acral lentiginous melanoma^32^.

Consequently, morphologic alterations associated with mutated *BRAF* are likely too subtle to be detected through traditional microscopy. Saliency maps or explainability techniques alter individual pixels and capture its effects on model performance^34^. In our study, we developed a novel pipeline that generates saliency maps for our network mutation prediction model and identified pixels corresponding to cellular nuclei as important for network decision-making (**Figure 6**) in both our institutional cohort as well as the TCGA cohort. We further investigate whether there are nuclear features that are associated with *BRAF* mutational status using pathomics to extract and quantitate 18 features using CellProfiler software^24^. Nuclei harboring mutated *BRAF* were larger and rounder than wild-type *BRAF* nuclei as measured by area, radii, and eccentricity. Notably, this corroborates previous studies that described *BRAF*-mutated melanomas as featuring larger and epithelioid melanocytes^32, 33^.

Because WSI analysis is a crucial feature for clinical adaptability, we also built a fully automated model that first applies a tumor selection algorithm (TumorNet) on non-annotated images followed by the mutation prediction algorithm. With the recent FDA approval of the first WSI imaging system for primary diagnosis in pathology^35^, the digitization of slides seems poised to be integrated into routine clinical practice. For instance, feature extraction from WSI analysis integrated with clinicopathologic data, mutational status, and gene expression data led to an improved prognostic model for recurrence-free survival in melanomas from TCGA^36^, while pathomics combined with trancriptomics analysis of CD8(+) T-cell distribution in metastatic melanomas can potentially predict clinical responses to BRAF-inhibitor therapy^37^. Other approaches have combined radiomics with pathomics to localize high-grade prostate cancers^38^ or predict for outcomes such as recurrence-free survival in lung cancer patients^39^.

Similarly, while deep learning-based mutational predictions are unlikely to replace direct molecular testing on tissue in the immediate future, there is great promise for these computational approaches to be integrated into higher order models, such as predicting for treatment responders vs. non-responders or survival outcomes, as has been previously demonstrated in lung cancers^40^ and gliomas^41^. We present a fully automated deep CNN model that accurately differentiates melanomas from benign tissue and uses morphologic features to predict the presence of the *BRAF* driver mutations on two independent cohorts. We confirm that the mutated *BRAF* genotype is linked to phenotypic alterations at the level of the nucleus through saliency mapping and pathomics analysis, providing additional insights on how this mutation affects tumor structural characteristics. Compared to direct testing methods, such an image-based approach has the potential to provide mutational data in a rapid, cost-reducing, and tissue-sparing manner that can be scaled up in research or even possibly, clinical settings.

## Materials and Methods

### Dataset of whole-slide images

All patients were enrolled in an IRB-approved clinicopathological database and biorepository in the Interdisciplinary Melanoma Cooperative Group (IMCG) at NYU Langone Health. The IMCG collects prospective clinical, pathological, and follow-up data from melanoma patients who present for diagnosis and/or treatment^42^.

365 H&E-stained FFPE whole-slides from 324 primary melanomas diagnosed between 1994 to 2013 were retrieved and digitized at 20x magnification. A single board-certified dermatopathologist (RHK) reviewed all digitized slides for image quality and excluded images that were blurry, faded, or did not contain any tumor. 293 images from 256 melanomas were subsequently annotated by RHK for tumor-rich regions of interest (ROIs) using Aperio ImageScope software. Driver mutations were previously determined by Sanger sequencing.

### Dataset from The Cancer Genome Atlas

68 FFPE slides of primary melanomas from 66 patients from the TCGA were downloaded and tiled into non-overlapping tiles of 299×299 pixels. Clinical information was not available for all slides. 28 slides with a Breslow depth similar to our cohort were maintained a second independent cohort. All tiles were sorted for testing and TFRecord files were generated. The slides were passed through the mutation prediction networks and the average probabilities per slide were used for the AUC calculation. The TCGA cohort was used as a generalizability metric for our classifiers. 64 slides were used for the pathomics analysis. The 4 slides excluded were very large and generated a very high number of tiles making the processing time for CellProfiler prohibitive.

### Software availability

We utilized the adapted Tensorflow DeepPATH pipeline (https://github.com/ncoudray/DeepPATH.git) to perform our analysis using the Inception v3 CNN architecture. To train the vgg16 and resnet18 architectures we used the PathCNN pipeline published on github (https://github.com/sedab/PathCNN). Our CellProfiler analysis pipeline is also available on github (https://github.com/sofnom/HistoPathNCA_pipeline).

### Image pre-processing

#### BRAF mutation prediction

To avoid introducing potential bias in our BRAF mutation classifiers, only the slide with the highest tumor content was used per patient, resulting in a dataset of 256 slides. WSI were partitioned at 20x magnification into non-overlapping 299×299 pixel tiles. For these classifiers, only the tiles from the area annotated as tumor were included in the analysis. This process generated 222,561 total tiles in our dataset, after removing tiles with more than 50% background (white area of slides). All tiles take the label of the slide they belong to and are sorted in training, validation and independent sets comprising of 70%, 15% and 15% of the total number of tiles correspondingly. All tiles from a specific slide are included in the same set with no overlap allowed. Tile sorting was performed using sorting option number 14 from the DeepPATH pipeline. Tiles in the train and validation sets were then converted to TF record format, which is necessary for training of Inception v3, in groups of 1024 tiles in each TF record file for the training set and 128 tiles for the validation set.

#### Tumor annotation network

All 293 whole-slide images were tiled for this task in order to provide the maximum amount of data available for training, similar to a data augmentation technique. The slides were tiled separately for the areas annotated as “tumor” and “non-tumor”. The number of tiles is presented in **Supplemental Table 1,** for all three magnifications explored. Tile sorting was performed using sorting option 19 from the DeepPATH pipeline. Tiles in the train and validation sets were converted as before to TF record format in groups of 1024 tiles in each TF record file for the training set and 128 tiles for the validation set.

### Deep learning with Convolutional Neural Networks for BRAF mutation prediction

#### Inception v3

The Inception v3 architecture is a Convolutional Neural Network (CNN) that utilizes modules comprised of various convolutions with different kernel sizes and a max pooling layers. The network was trained on 70% of the tiles from each data set, with 15% of the tiles used for validation and 15% used for independent testing.

The network was trained from scratch and using transfer learning for 150,000 training steps on batches of 160 images, on 4 GPUs. The corresponding number of epochs varies based on the total number of tiles and the batch size and is determined by the following equation:

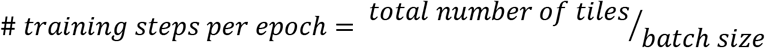

The learning rate was set to 0.1. For transfer learning, the initial learning rate was set to 0.001. The RMSProp for gradient descent optimizer was used with learning rate decay factor of 0.16 and 15 epochs per decay for both training modes. The activation function used in the output layer was softmax. The built-in data augmentation techniques of Inception v3 were utilized as defined in the “image_processing.py” script available here https://github.com/ncoudray/DeepPATH/tree/master/DeepPATH_code/01_training/xClasses/inception. These include horizontal flip of the images and random color distortion, as well as obtaining randomly sized crops of the training images and resizing them to the necessary tile size.

For transfer training using ImageNet weights we used the checkpoint at the following link: http://download.tensorflow.org/models/image/imagenet/inception-v3-2016-03-01.tar.gz. For transfer training using the weights from the corresponding tumor annotation classifier we used the best checkpoints of the tumor classification networks.

The network’s performance was monitored based on the AUC on the validation set. The best performing model was chosen either when the validation AUC displayed a very sharp drop between the training steps or when there was a clear plateau. The performance of the best model was then evaluated on the independent set and the AUC was calculated. The network outputs a probability value for every tile for each class of interest. The tile is assigned to the class with the highest probability. The tile probabilities are then averaged to produce the final slide probability.

A heat map for each slide in the test set can be generated according to the “0f_HeatMap_nClasses.py” script in (https://github.com/ncoudray/DeepPATH.git). The heat map overlaps the probability information for each tile with the initial H&E slide to produce a color-coded image visualizing the localization of the mutation as predicted by the network at the tile level. The color intensity is analogous to the probability value of the tile to belong in each class.

#### VGG16 and ResNet18

These architectures were trained using the code available at https://github.com/sedab/PathCNN. They were trained for 50 training epochs, using learning rate of 0.1 for VGG16 and 0.05 for ResNet18. Image tiles are automatically resized from 299×299 to the default tile size for these architectures which is 224×224 pixels. The SGD optimizer is used. Dropout was set at 0.1 and the Xavier initialization was employed. Data augmentation included random horizontal image flip, random image rotation and random color normalization, as defined in the “train.py” script of the pipeline. A leaky non-linear function was used. Network performance was measured by the AUC on the validation set. The best checkpoint was chosen the same way as for the Inception v3 architecture above.

#### Hardware

All deep learning models were trained on Tesla V100-SXM2-16G GPUs.

#### Automated tumor selection classifiers - TumorNet

Inception v3 was trained from scratch on the tiles generated as described under “Image preprocessing, Tumor annotation network” on 4 GPUs. Learning rate was set to 0.1 and batch size to 400, for all magnifications. Softmax was used as the activation function for the output layer. Training loss and validation AUC were monitored the same way as for the BRAF mutation classifiers and performance is measured on the independent test set for the best model.

#### BRAF mutation prediction on the automatically selected tumor areas for the sequential model

To be able to use a sequential model with automatic tumor annotation we wanted to show that a BRAF prediction classifier trained on automatically selected tumor regions will achieve similar AUC as the one trained on the manually selected ones. We passed all 256 slides through the TumorNet20x network to annotate the tumor regions. We then split the selected tumor tiles (tumor probability >= 0.365789) into training, validation and test sets. Inception v3 was trained on the tiles that are considered as tumor, using transfer training on the best TumorNet checkpoint at 20x magnification with the same parameters as for the manually annotated tumor regions.

### Statistical analysis

After training and choosing the best performing model on the validation set, model performance was evaluated using the independent test set, which is comprised of a held-out population of tiles coming from 36 slides. Each slide comes from a unique patient in the case of our BRAF prediction classifiers. Regarding the tumor annotation classifiers, where each patient can have multiple slides, we report the “per patient” AUC. The probabilities for each slide were aggregated by the average of probabilities of the corresponding tiles. Receiver Operative Characteristic (ROC) curves and the corresponding Area Under the Curve (AUC) were generated as a measure of accuracy. Heat maps allowed visualization of probability differences and regions of interest.

### Conventional Machine Learning Models for Pathomics

The multivariate logistic regression model was built using the *glm* function in R from the “ROCR” package. The Random Forest model was created using the *randomForest* function from the “randomForest” package in R.

### Smoothed training loss and validation AUC plots

Smoothing was performed using the function geom_smooth() from the ggplot2 package with default parameters based on the number of data points, in R.

### Receiver Operating Characteristic Curves

ROC curves were generated using the pROC package in R.

### Immunohistochemical analysis of mutated BRAF V600E

Immunohistochemistry (IHC) was performed on 10% neutral buffered FFPE, 4-μm human archival melanoma sample sections collected on plus slides (Fisher Scientific, Cat# 22-042-924) and stored at room temperature. Unconjugated, mouse anti-human Serine-Threonine-Protein Kinase B-raf (BRAF) V600E, clone VE1 (Abcam Cat# ab228461, Lot# GR32335840-6) raised against a synthetic peptide within human BRAF (amino acids 550-650) containing the glutamic acid substitution, was used for IHC^43, 44^. BRAF antibody was optimized on known positive and negative colon samples and subsequently validated on a mixed set 20 known positive/negative samples. Chromogenic immunohistochemistry was performed on a Ventana Medical Systems Discovery Ultra using Ventana’s reagents and detection kits unless otherwise noted. In brief, slides were deparaffinized online and antigen retrieved for 24 minutes at 95°C using Cell Conditioner 1 (Tris-Borate-EDTA pH8.5). BRAF was diluted 1:50 in Ventana antibody diluent (Ventana Medical Systems, Cat# 251-018) and incubated for 16 minutes at 36°C. Endogenous peroxidase activity was post-primary blocked with 3% hydrogen peroxide for 4 minutes. Primary antibody was detected using Optiview linker followed by multimer-HRP incubated for 8 minutes each, respectively. The complex was visualized with 3,3 diaminobenzidene for 8 minutes and enhanced with copper sulfate for 4 minutes. Slides were counterstained online with hematoxylin for 8 minutes and blued for 4 minutes. Slides were washed in distilled water, dehydrated and mounted with permanent media. Positive and negative (diluent only) controls were run in parallel with study sections. Blinded analysis of staining was performed by a single dermatopathologist (GJ).

### BRAF V600E-predicted tumor areas overlap with immunohistochemical V600E antibody staining

Manual annotation of V600E-stained areas on the IHC slides was performed using the Aperio ImageScope software. The same area was annotated on the H&E slide by visual overlap of the slides by a single certified dermatopathologist (GJ). Different tumor slices are used for IHC and H&E and most available alignment software are not allowing for image rotation which would account for a more faithful image alignment. Consequently, they were deemed unreliable to overlap the stained regions with the tumor area of the H&E slide. Manual annotation is more reliable in this case. After obtaining the desired masks, the probability distributions for tiles assigned to the V600E-stained areas as opposed to the probabilities of the remaining tumor tiles were plotted in the form of a boxplot for all 17 BRAF V600E slides. P-values were calculated using an unpaired two-sided Wilcoxon rank sum test for each slide.

### Generating saliency maps

Saliency maps were created with the Smooth Integrated Gradients method^45^. First, an InceptionV3-architectured graph was constructed using Tensorflow slim API in order to reload the trained model. The architecture and all hyperparameters were kept exactly the same as the trained model. Then, selected tiles from the independent test set and the TCGA cohort with the highest and the lowest predicted probabilities were fed into the reloaded models. The weights of the layer before the last fully connected layer were then used to build the saliency map. We used the Saliency package (https://pypi.org/project/saliency/) from PyPI to generate Smoothed Integrated Gradients for these tiles. Considering the nature of the digital histopathology images, both pure black (RGB=[0,0,0]) and pure white (RGB=[255,255,255]) were used as the baselines to calculate the gradients. Saliency maps using the white background are presented in **Figure 6**. A better visualization output was made by overlaying saliency maps onto the original tiles.

### Pathomics Analysis using CellProfiler

To perform Pathomics analysis we used CellProfiler^24^, a publicly available software platform for cell and nuclear analysis from multiple formats of biological images. We developed a pipeline on CellProfiler version 3.1.8 to measure nuclear and cellular features on the tile level of H&E slides. CellProfiler 3 documentation is available here http://cellprofiler-manual.s3.amazonaws.com/CellProfiler-3.0.0/index.html for a detailed description of all pipeline steps that follows.

### Pipeline steps for nuclear annotation

#### UnmixColors

The pipeline starts by de-convolving the Hematoxylin, Eosin and Pigment signals and generating grayscale images indicating the location of each stain with white color. The deconvolution of Hematoxylin and Eosin is built-in the software and the pigment color was determined by choosing a custom color profile based on the pigmentation of our images.

#### IdentifyPrimaryObjects

Then, the Hematoxylin stain is used to annotate nuclei, and the pigment stain is used to annotate pigmented regions on the tile. To annotate nuclei, we decided to adopt the Otsu method with default parameters except for “threshold correction factor” for which we used value of 1.3 instead of the default 1.0 for more stringent annotation. “Typical diameter of objects” was set to 10 to 40 pixels, as by default. For pigment annotation, we used a manual thresholding method with a threshold of 0.8 and ‘typical diameter of objects” was set to 10 to 100 to reduce the number of objects identified. Our slides were not color normalized. We noticed that color normalization was interfering with the annotation of pigmented regions because it was reducing the contrast between the pigment color and the rest of the slide. Instead, we opted for the Otsu method which tests multiple thresholding values before performing nuclear annotation, therefore it automatically adapts to each tile’s color profile. For cell annotation, we changed the annotation method to Minimum Cross Entropy with the default thresholding smoothing value of 1.3488 and the default threshold correction factor of 1.0. The rest of pipeline stages are unchanged.

#### ConvertObjectsToImage

This step is used to convert the identified pigment objects to a mask image that can be used by the following step *MaskObjects.*

#### MaskObjects

Pigmented areas were excluded from our nuclear annotation because pigmented cells may represent melanophages rather than tumor cells.

#### OverlayOutlines

This step is overlaying the tile image with the identified nuclei and pigmented regions for visualization and evaluation of our pipeline. The objects are overlayed using the default parameters.

#### SaveImages

The overlay images of can be saved in a jpeg format.

#### MeasureObjectSizeShape

This module measures object size and shape features. In total, it measures 18 features:

#### Export ToSpeadsheet

This step is used to save the outputs of the previous step into a text file for every slide.

Our code is available on github: https://github.com/sofnom/HistoPathNCA_pipeline.

### Processing of CellProfiler results

The CellProfiler pipeline generates data for each tile of all slides of a patient. All identified cells and nuclei per patient were collected and the nuclear and cellular features were averaged by patient. Additional normalization to the total number of tiles by patient was needed for the total number of objects, total object area and total pigmented area. The distribution of each feature was plotted for both the non-tumor and the tumor tiles, stratified by the true label of the patient. Logarithmic conversion was used for plotting the total object and pigment areas. P-values for the boxplots were calculated using an unpaired two-sided Wilcoxon rank sum test in R.

### Down-sampled training of Inception v3

The NYU dataset was down-sampled to 20, 40, 60 and 80% of the available slides. We made sure to maintain the same proportion of BRAF mutant to BRAF WT slides in the down-sampled datasets as for the network trained on the initial dataset to avoid biasing the training process and our results (**Supplemental Table 4**). Transfer training at 20x magnification was performed using the TumorNet weights. Learning rate was set to 0.1 and batch size to 160. All other training parameters were the same as the network trained on the whole dataset. The average AUC on the validation and test sets was calculated for the best checkpoints along with the average CIs. The data were imported in Microsoft Excel. Using the built-in “Power” function we fit an inverse power law curve to the data to predict the number of available tiles we would need to achieve a BRAF mutation prediction AUC of 80% and 90%; performance which is much more relevant for clinical practice.

## Supplementary Materials

**Fig S1.** Training of tumor annotation classifier at multiple magnifications.

**Table S1.** Training Tumor Annotation Network for different magnifications.

**Table S2.** Training multiple architectures for BRAF mutation prediction.

**Fig S2.** Different learning modes affect *BRAF* mutation prediction (Inception v3; 20x magnification).

**Fig S3.** Different learning modes affect *BRAF* mutation prediction (Inception v3; 10x magnification).

**Fig S4.** Different learning modes affect *BRAF* mutation prediction (Inception v3; 5x magnification).

**Table S3.** Different learning modes affect *BRAF* mutation prediction (Inception v3).

**Table S4.** Down-sampled datasets for Inception v3 training.

**Fig S5.** Dataset down-sampling reduces classifier’s performance.

**Fig S6.** BRAF mutation prediction using manual vs. network annotated tumor areas.

**Fig S7.** Nuclear features for NYU cohort.

**Fig S8.** Nuclear features for TCGA cohort.

**Fig S9.** Cellular features for NYU cohort.

**Fig S10.** Cellular features for TCGA cohort.

**Table S5.** Pathomics machine learning models for BRAF mutation prediction using nuclear features.

## Acknowledgments

We thank Luis Chiriboga from the NYU Experimental Pathology Immunohistochemistry Core Laboratory. The results shown here are in part based upon data generated by the TCGA Research Network: https://www.cancer.gov/tcga. This work has used computing resources at the High-Performance Computing Facility at the NYU Medical Center. We also want to thank Anna Yeaton for discussions about this project.

## Author contributions

Study concept and design: RHK, SN, IO, AT. Acquisition of data: RHK, SN, ZD, GJ, UM, RLS, RSB. Analysis and interpretation of data: RHK, SN, NC, GJ, JSW, NR, IO, AT, RH, EE; IA and DF offered constructive feedback and suggestions. Study supervision: NC, IO, AT.

