## Supplemental Material for "A Deep Learning Approach for Rapid Mutational Screening in Melanoma"

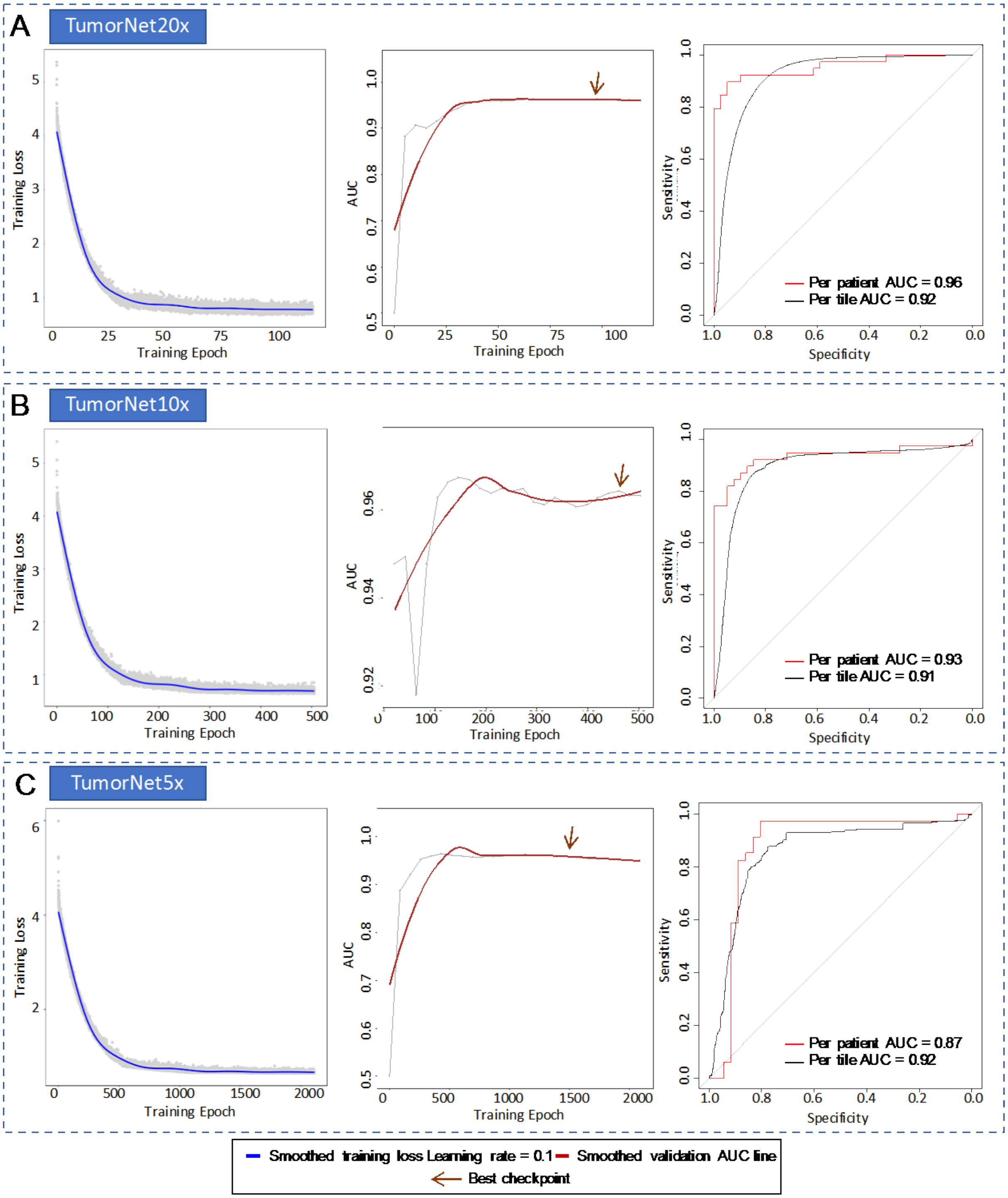


**Supplemental Figure 1**. **Training of tumor annotation classifier at multiple magnifications**. **A**. **20x** Training loss curve (left). AUC on validation set. Best checkpoint is chosen at 95k training steps (middle). ROC curve on test set for best checkpoint (right). **B. 10x.** Training loss curve (left). AUC on validation set. Best checkpoint is chosen at 110k training steps (middle). ROC curve on test set for best checkpoint (right). **C. 5x.** Training loss curve (left). AUC on validation set. Best checkpoint is chosen at 85k training steps (middle). ROC curves on test set for best checkpoint (right). Network performance is similar for all three magnifications. The slightly reduced performance for 5x can be explained by the smaller number of available tiles for training.

**Supplemental Table 1. Training Tumor Annotation Network for different magnifications.**

| Magnification | Tumor tiles | Non-tumor tiles | Per patient Test AUC | Per Tile Test AUC |
|---|---|---|---|---|
| 20x | 252,372 | 351,920 | 0.96 [95% CI:0.90-0.99] | 0.919 [95% CI:0.918-0.921] |
| 10x | 62,307 | 81,241 | 0.93 [95% CI:0.86-0.99] | 0.908 [95% CI:0.903-0.912] |
| 5x | 15,126 | 18,289 | 0.87 [95% CI:0.76-0.96] | 0.921 [95% CI:0.912-0.958] |

**Supplemental Table 2. Training multiple architectures for BRAF mutation prediction.**

| Architecture | Per patient Test AUC | Per patient TCGA AUC |
|---|---|---|
| Inception v3 | 0.69 [95% CI:0.50-0.86] | 0.73 [95% CI:0.53-0.94] |
| Vgg16 | 0.74 [95% CI:0.58-0.90] | 0.59 [95% CI:0.37-0.82] |
| Resnet18 | 0.86 [95% CI:0.74,0.99] | 0.59 [95% CI:0.36-0.81] |

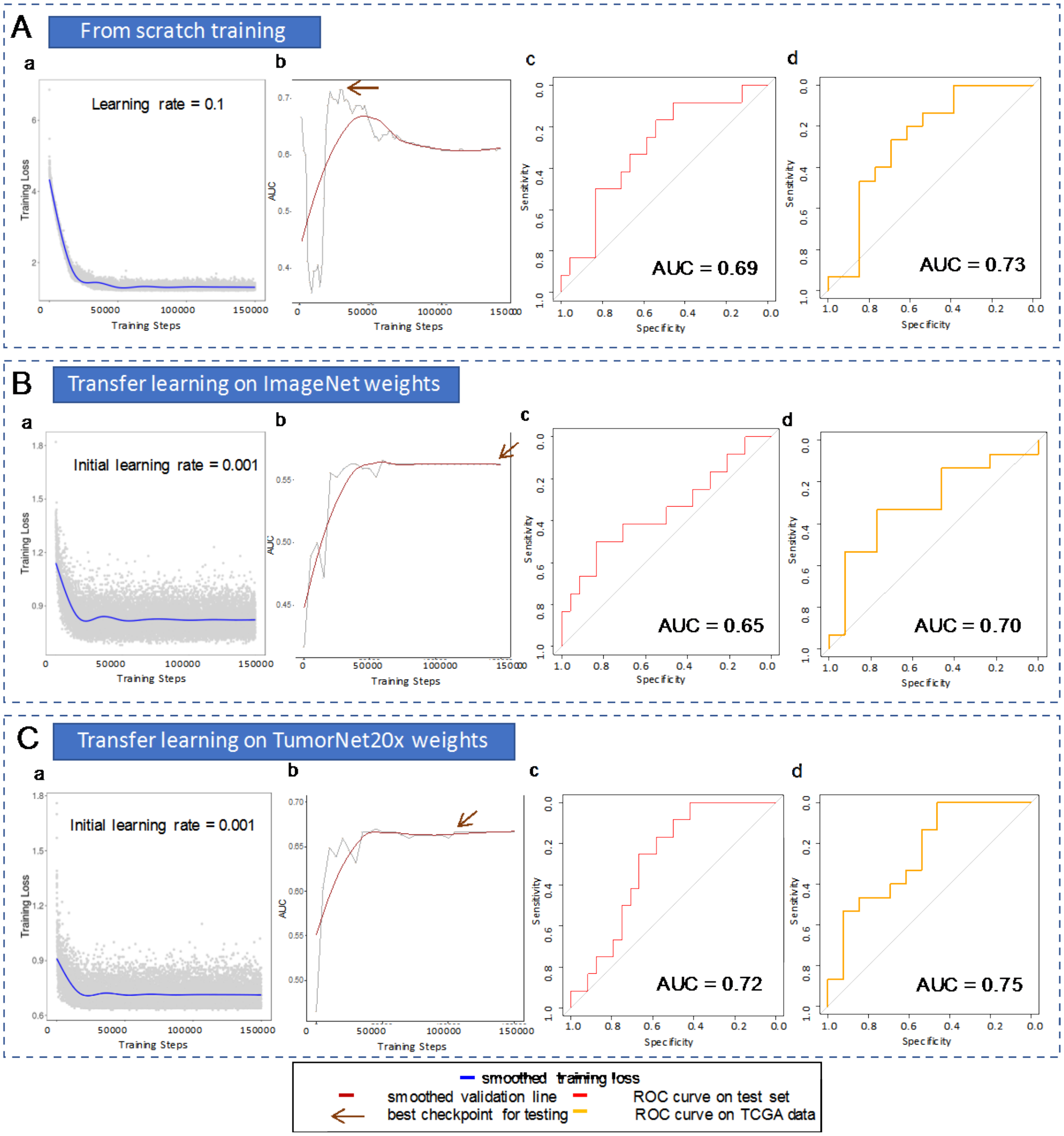


**Supplemental Figure 2. Different learning modes affect *BRAF* mutation prediction (Inception v3; 20x magnification). A. Training from scratch**. **a)** Training Loss curve for training from scratch. **b)** Validation AUC across training. Best checkpoint is chosen at 31,500k training iterations. **c)** ROC curve for independent test set on best checkpoint. AUC is 0.69. **d)** ROC curve for external TCGA cohort on best checkpoint. AUC is 0.73. **B. Transfer training on ImageNet weights. a)** Training Loss. **b)** Validation AUC across training. Best checkpoint is chosen at 150k training iterations. **c)** ROC curve for independent test set on best checkpoint. AUC is 0.65. **d)** ROC curve for external TCGA cohort on best checkpoint. AUC is 0.70. **C. Transfer training on the weights from the 20x tumor annotation network**. **a)** Training Loss. **b)** Validation AUC across training. Best checkpoint is chosen at 95k training iterations. **c)** ROC curve for independent test set on best checkpoint. AUC is 0.72. **d)** ROC curve for external TCGA cohort on best checkpoint. AUC is 0.75.

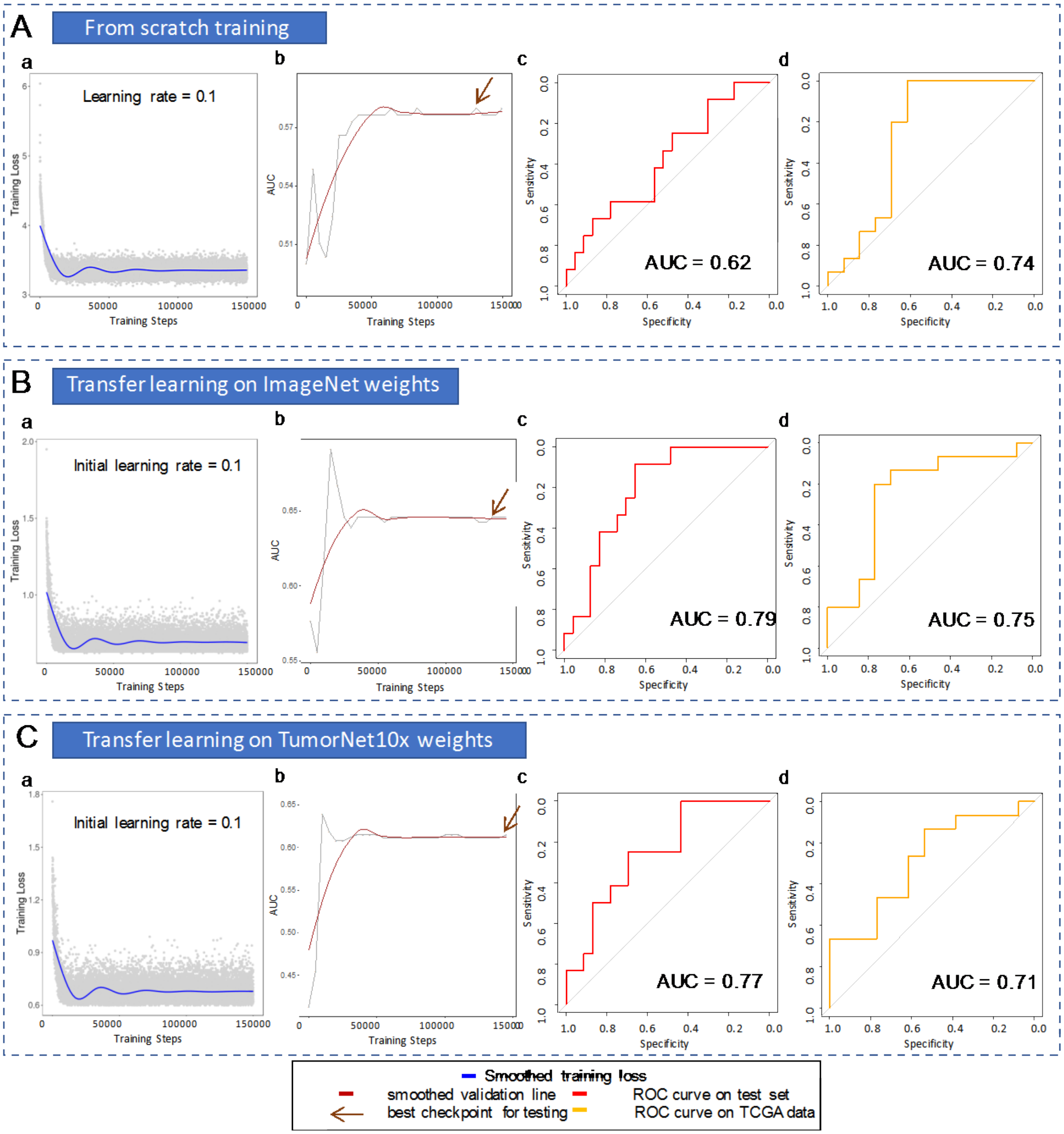


**Supplemental Figure 3. Different learning modes affect *BRAF* mutation prediction (Inception v3; 10x magnification). A. Training from scratch**. **a**) Training Loss of training from scratch. **b)** Validation AUC across training. Best checkpoint is chosen at 130k training iterations. **c)** ROC curve for independent test set on best checkpoint. AUC is 0.62. **d)** ROC curve for external TCGA cohort on best checkpoint. AUC is 0.74. **B. Transfer training on ImageNet weights**. **a)** Training Loss. **b)** Validation AUC across training. Best checkpoint is chosen at 140k training iterations. **c)** ROC curve for independent test set on best checkpoint. AUC is 0.79. **d)** ROC curve for external TCGA cohort on best checkpoint. AUC is 0.75. **C. Transfer training on the weights from the 10x tumor annotation network**. **a)** Training Loss. **b)** Validation AUC across training. Best checkpoint is chosen at 145k training iterations. **c)** ROC curve for independent test set on best checkpoint. AUC is 0.77. **d)** ROC curve for external TCGA cohort on best checkpoint. AUC is 0.71.

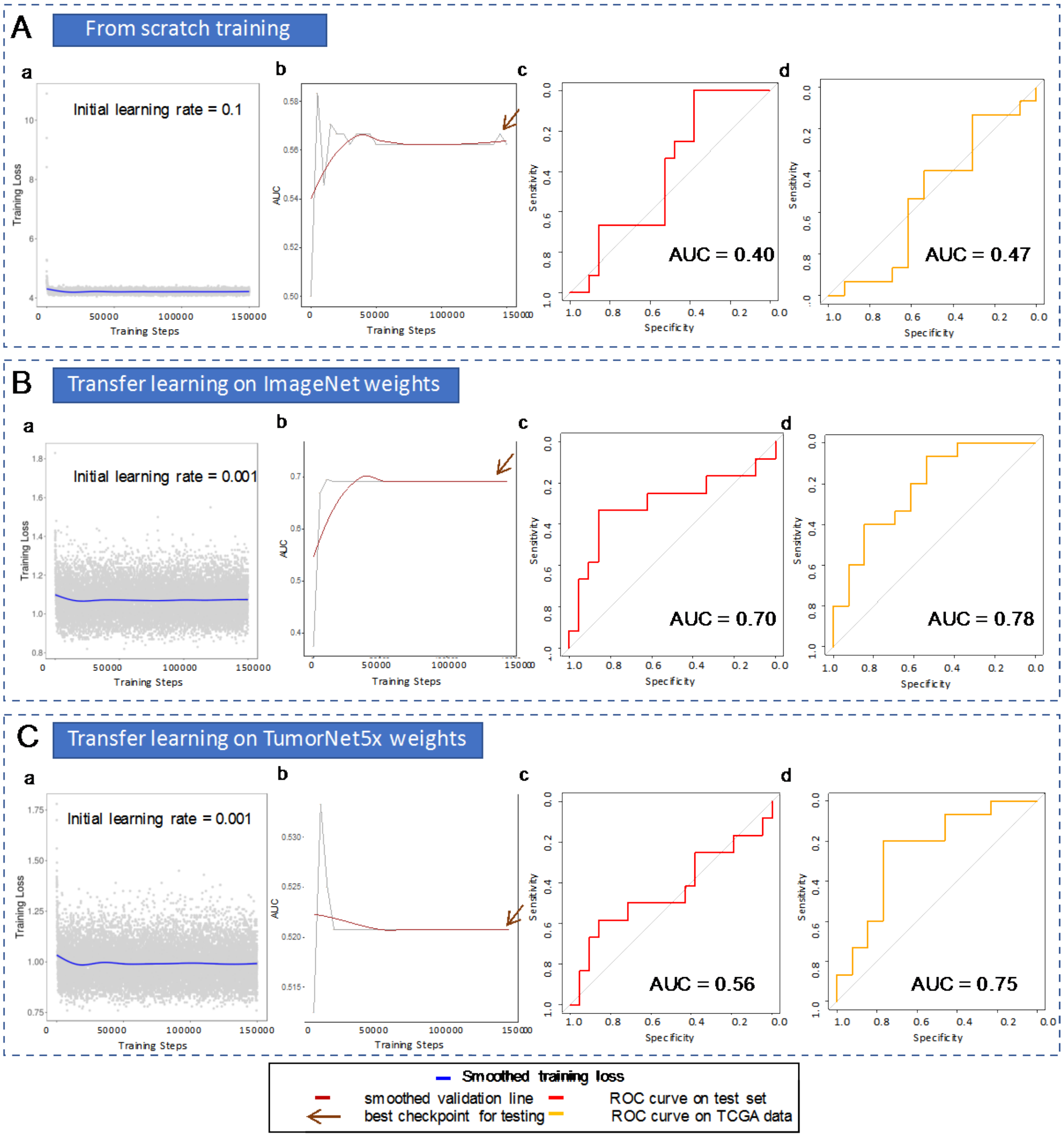


**Supplemental Figure 4**. **Different learning modes affect *BRAF* mutation prediction (Inception v3; 5x magnification). A. Training from scratch**. a) Training Loss of training from scratch. **b)** Validation AUC across training. Best checkpoint is chosen at 145k training iterations. **c)** ROC curve for independent test set on best checkpoint. AUC is 0.40. **d)** ROC curve for external TCGA cohort on best checkpoint. AUC is 0.47. **B. Transfer training on ImageNet weights**. **a)** Training Loss. **b)** Validation AUC across training. Best checkpoint is chosen at 145k training iterations. **c)** ROC curve for independent test set on best checkpoint. AUC is 0.70. **d)** ROC curve for external TCGA cohort on best checkpoint. AUC is 0.78. **C. Transfer training on the weights from the 5x tumor annotation network. a)** Training Loss. **b)** Validation AUC across training. Best checkpoint is chosen at 145k training iterations. **c)** ROC curve for independent test set on best checkpoint. AUC is 0.56. **d)** ROC curve for external TCGA cohort on best checkpoint. AUC is 0.75.

| **Supplemental Table 3. Different learning modes affect *BRAF* mutation prediction (Inception v3).** | | | |
|---|---|---|---|
| **Magnification** | **Training mode** | **Test AUC** | **TCGA AUC** |
| 20x | scratch | 0.69 [95% CI:0.50-0.86] | 0.73 [95% CI:0.53-0.94] |
| 20x | ImageNet | 0.65 [95% CI:0.44-0.85] | 0.70 [95% CI:0.49-0.90] |
| 20x | TumorNet | 0.72 [95% CI:0.53-0.87] | 0.75 [95% CI:0.57-0.94] |
| 10x | scratch | 0.62 [95% CI:0.42-0.81] | 0.74 [95% CI:0.52-0.95] |
| 10x | ImageNet | 0.79 [95% CI:0.63-0.93] | 0.75 [95% CI:0.55-0.95] |
| 10x | TumorNet | 0.77 [95% CI:0.61-0.91] | 0.71 [95% CI:0.52-0.91] |
| 5x | scratch | 0.40 [95% CI:0.22-0.69] | 0.47 [95% CI:0.24-0.70] |
| 5x | ImageNet | 0.70 [95% CI:0.47-0.91] | 0.78 [95% CI:0.60-0.96] |
| 5x | TumorNet | 0.56 [95% CI:0.32-0.77] | 0.75 [95% CI:0.56-0.95] |

| **Supplemental Table 4. Down-sampled datasets for Inception v3 training.** | | | |
|---|---|---|---|
| | **Number of slides per mutation in down-sampled dataset** | | |
| **Percentage** | **BRAF mutant** | **BRAF WT** | **Total** |
| **20%** | 17 | 34 | 51 |
| **40%** | 34 | 68 | 102 |
| **60%** | 51 | 102 | 153 |
| **80%** | 68 | 136 | 204 |
| **100%** | 85 | 171 | 256 |

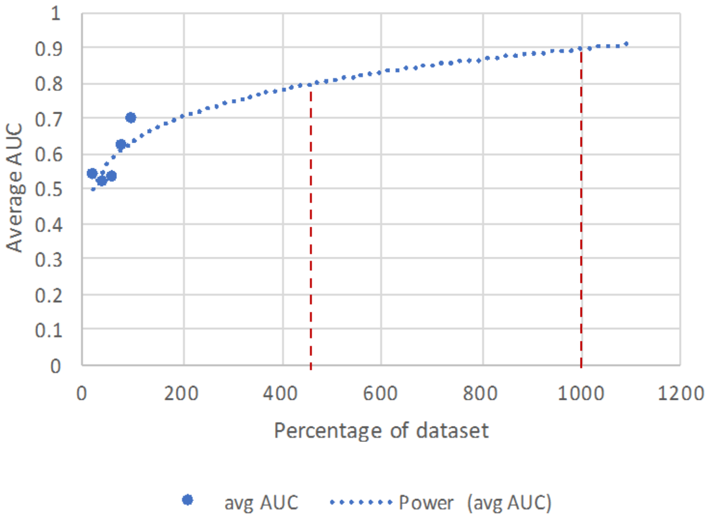


| percentage | avg AUC across validation and test sets | avg Low CI | avg High CI |
|---|---|---|---|
| 20 | 0.54 | 0.1 | 0.93 |
| 40 | 0.52 | 0.235 | 0.795 |
| 60 | 0.53 | 0.28 | 0.77 |
| 80 | 0.62 | 0.395 | 0.815 |
| 100 | 0.70 | 0.505 | 0.855 |

**Supplemental Figure 5. Dataset down-sampling reduces classifier's performance.** Inception v3 was trained on 20,40,60 and 80% of slides in the initial dataset using transfer training and the weights of TumorNet20x. Average AUC on validation and test set was reduced as expected. An inverse power law curve was fitted to the data. To achieve AUC of 0.8 we would need ~4.5x more data and to achieve an AUC of 0.9, ~10x more data would be needed.

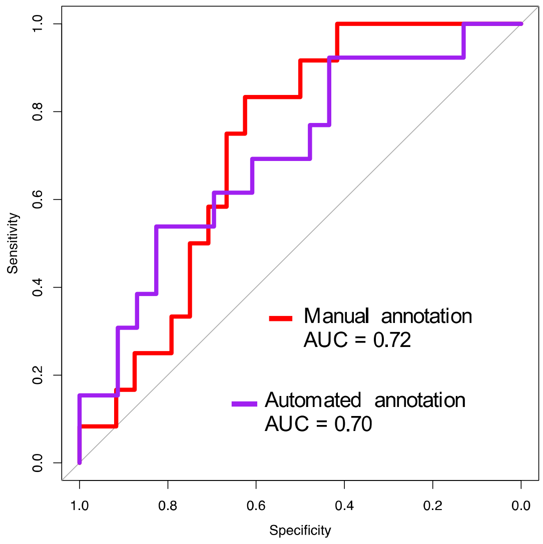


**Supplemental Figure 6**. **BRAF mutation prediction using manual vs. network annotated tumor areas.** BRAF classifier trained on the manually selected tumor regions achieved an AUC of 0.72 [95% CI: 0.54-0.87] and a BRAF classifier trained on the tumor regions automatically annotated by our tumor annotation network (TumorNet) achieved a similar AUC of 0.70, [95% CI: 0.54-0.87]

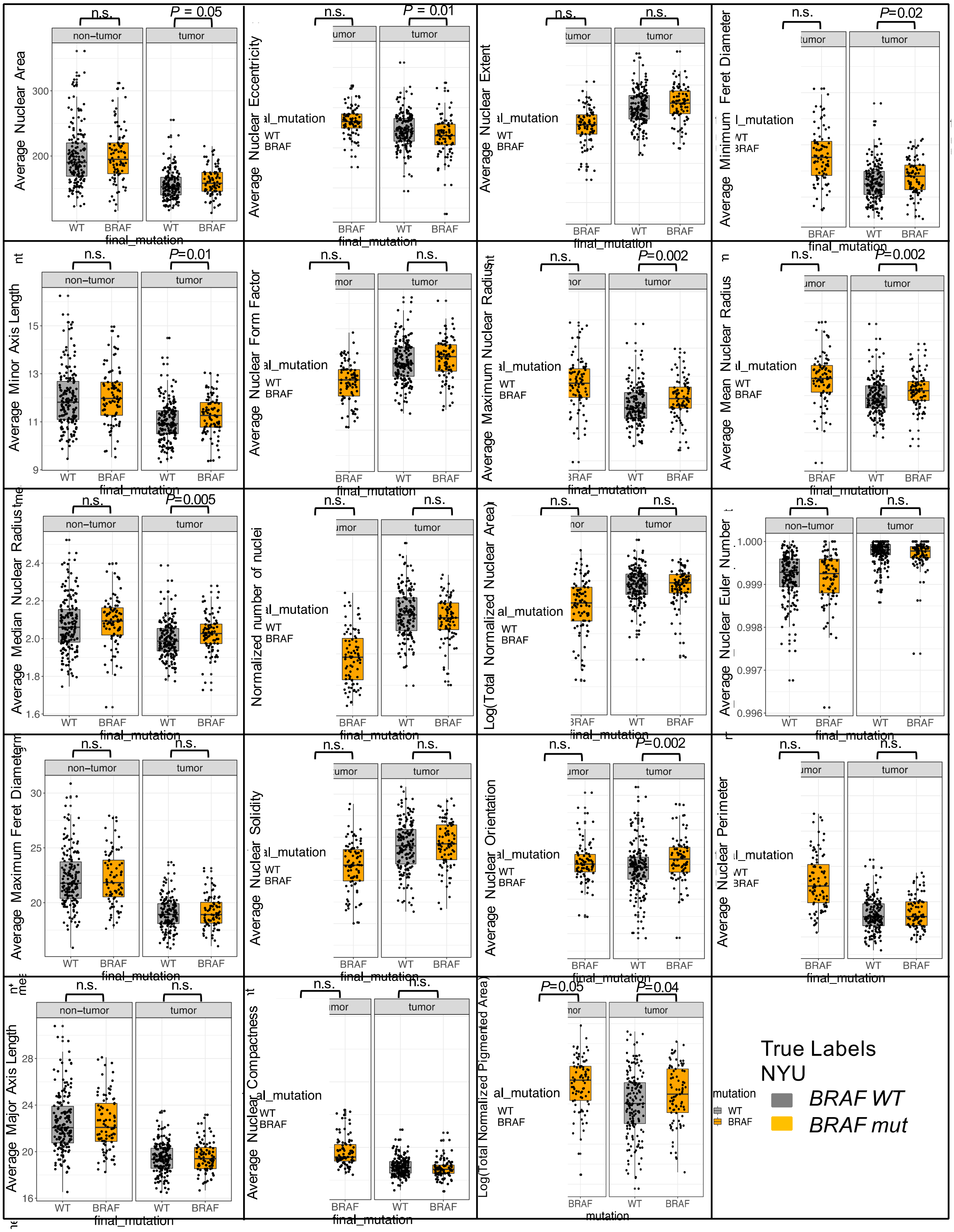


**Supplemental Figure 7.** Nuclear features for NYU cohort.

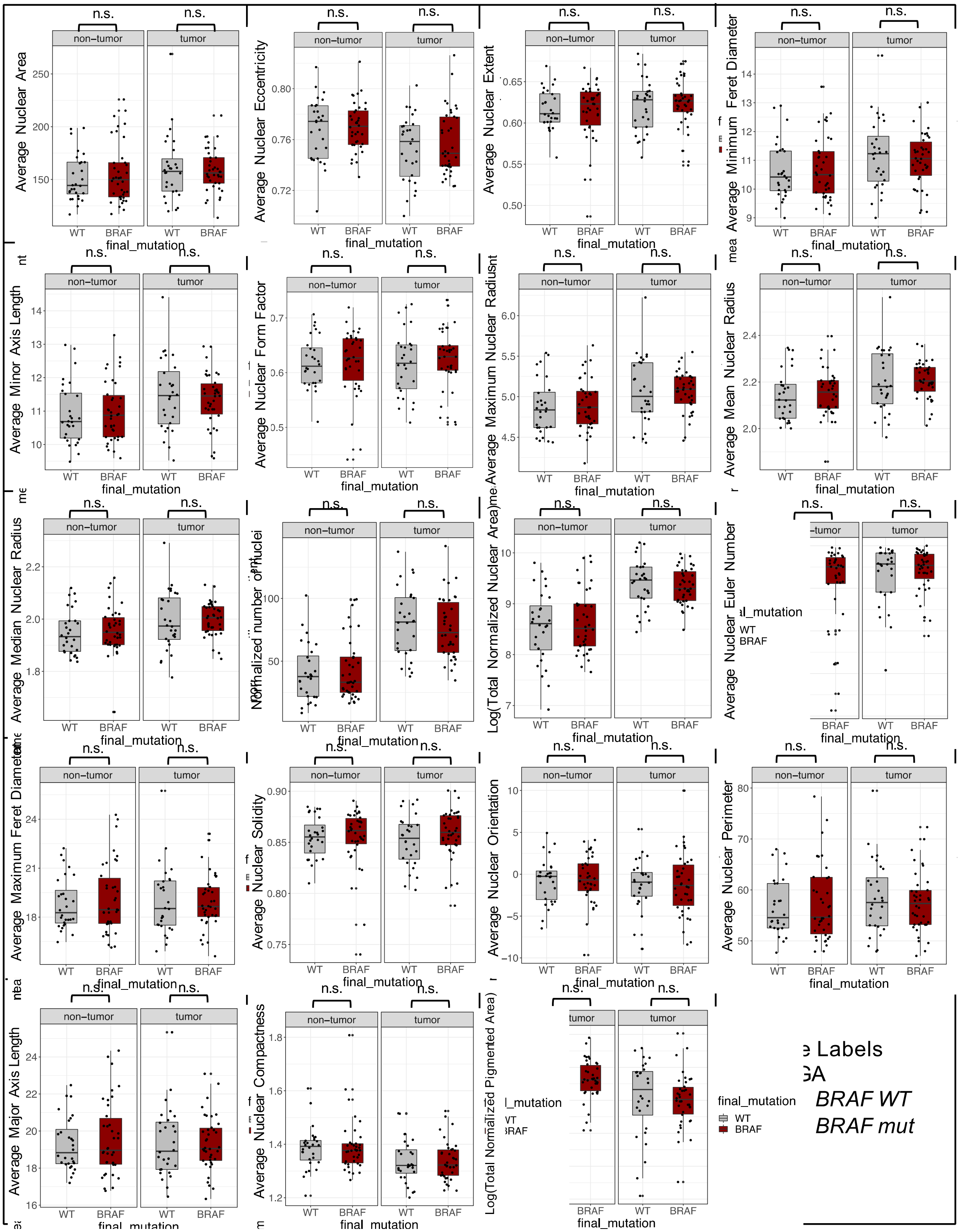

**Supplemental Figure 8.** Nuclear features for TCGA cohort.

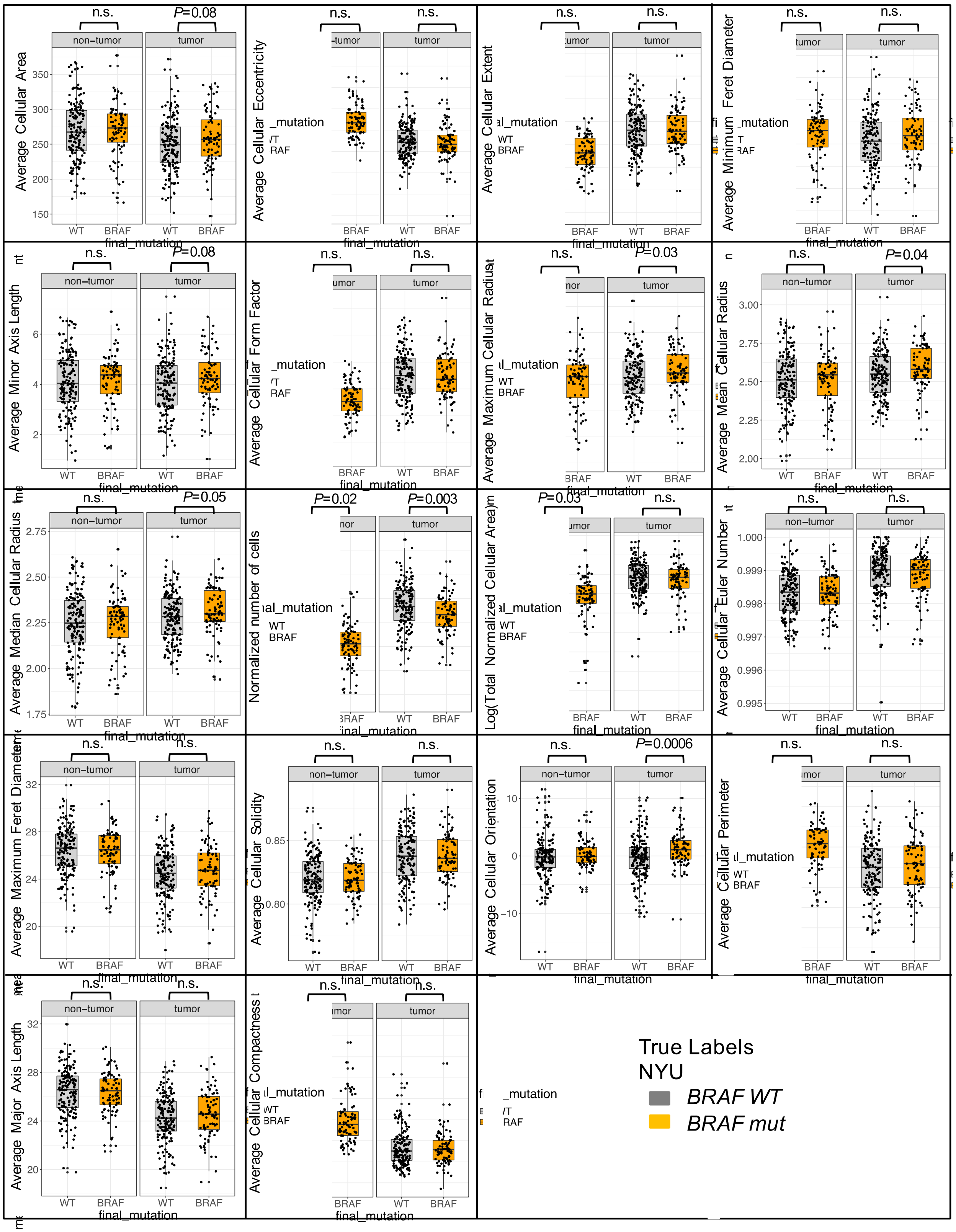

**Supplemental Figure 9.** Cellular features for NYU cohort.

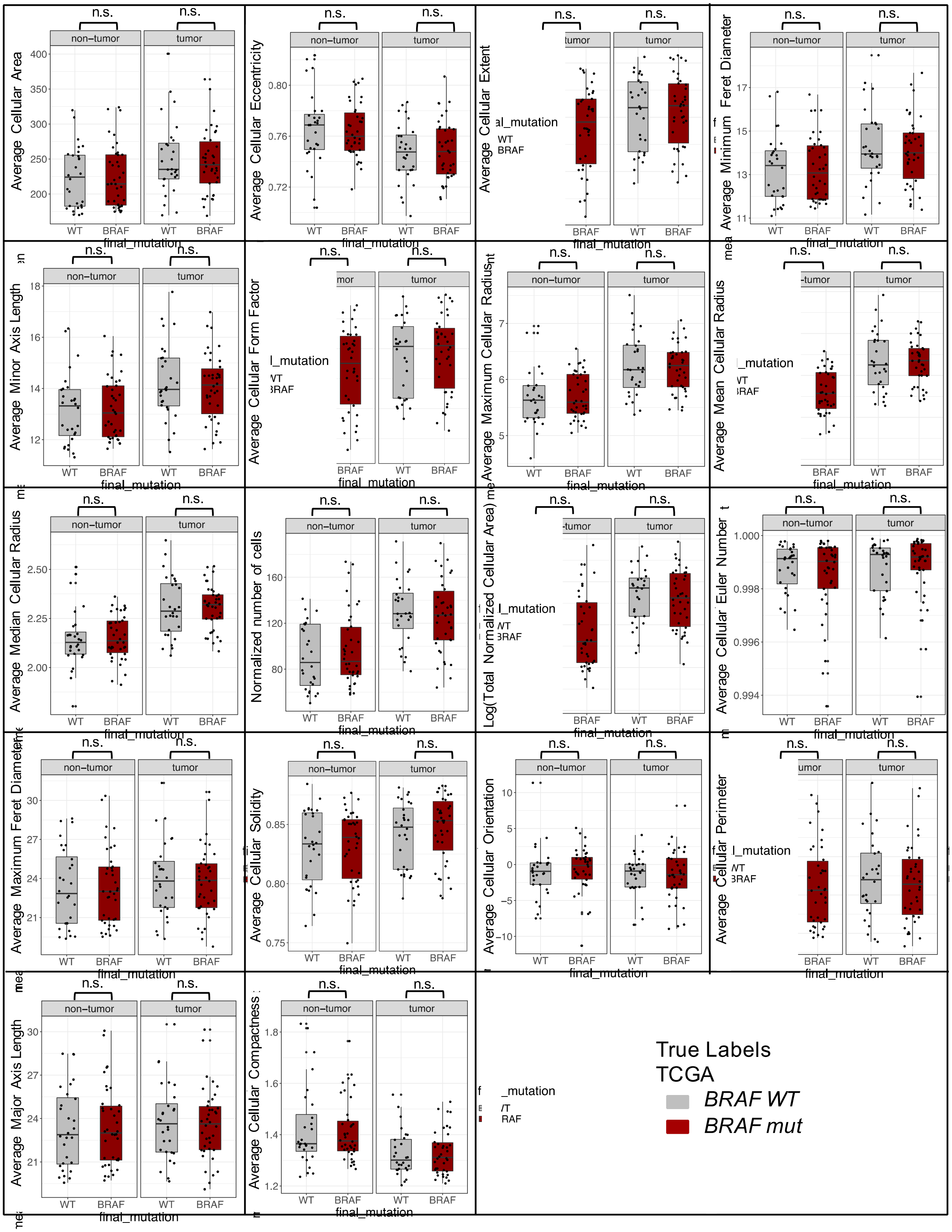


**Supplemental Figure 10.** Cellular features for TCGA cohort.

**Supplemental Table 5. Pathomics machine learning models for BRAF mutation prediction using nuclear features.**

| Model | 7-fold cross validation AUC on NYU dataset | 7-fold cross validation AUC on TCGA dataset |
|---|---|---|
| glm | 0.56, st.dev = 0.14 | 0.56, st.dev = 0.03 |
| Random forest | 0.58, st.dev = 0.11 | 0.61, st.dev = 0.02 |
